# Ancient environmental DNA reveals the distinct fingerprints of whaling and climate change on Antarctic marine biodiversity

**DOI:** 10.64898/2026.10.30.763115

**Authors:** Luke E. Holman, Matthew Mason, Lene Bruhn Pedersen, Juliane Müller, Frank Lamy, Evgeny A. Genelt-Yanovskiy, Giulia Zampirolo, Tobias Guldberg Frøslev, Robert J. W. Brewin, Ruth H. Thurstan, Callum M. Roberts, Mikkel Winther Pedersen, Shyam Gopalakrishnan, James Scourse, Kristine Bohmann

## Abstract

The Southern Ocean is a disproportionately large contributor to Earth system processes, playing a crucial role in the global transport of carbon, oxygen and heat. It also hosts enormous seasonal biological productivity that supports diverse endemic marine biodiversity. Industrial whaling in the first half of the 20th century and recent human-driven climate change have both profoundly altered Antarctic ecosystems, but ecological surveys of these impacts are limited to well-known taxa in the short period since monitoring began. Here, we analysed five sediment records to reconstruct more than two centuries of ecosystem change across the Western Antarctic Peninsula, spanning periods before and during major human impacts. Environmental DNA metabarcoding of these records revealed substantial biodiversity shifts, with change accelerating in recent decades. We find a significant role for both climate change and whaling in restructuring communities, identifying unique variance in whole-community change explained by each driver. Moreover, we find a negative association between whaling and metazoan families which feed from the water column, suggesting that pre-whaling ecosystems had greater pelagic productivity. Overall, our results reveal baseline Antarctic biodiversity from long before observational records and identify a distinct fingerprint of human drivers in this fragile, rapidly changing ecosystem.

## Introduction

The Southern Ocean is a crucial component of the Earth system. It hosts key up- and downwelling limbs of the global overturning circulation, which link ocean basins and connect the surface ocean to the deep sea^1,2^. This overturning supports an enormous flux of heat, carbon, oxygen and nutrients, giving the Southern Ocean a disproportionately large role in the climate system. Across the historical period it has absorbed an estimated 83% of heat from anthropogenic greenhouse gases and 43% of carbon emitted by human activities^3,4^. It is also a dominant site of ocean ventilation, accounting for about a third of oxygen supplied to the global ocean interior^5^.

These fluxes also sustain productivity well beyond Antarctic seas, with up to three-quarters of low-latitude primary production fuelled by nutrients originating in the Southern Ocean^6^. The region’s phytoplankton drive a biological carbon export to the deep sea that the rest of the ocean cannot fully compensate for if lost^7^, and this role extends up the food web, with substantial carbon export to the deep also supplied by large populations of zooplankton such as krill and salps^8,9^. High productivity means the Antarctic also supports many endemic species and enormous populations of migratory taxa. For example, the Southern Ocean hosts several million penguin breeding pairs across eight species^10–12^ and is a globally important feeding ground for migratory baleen whales, which travel thousands of kilometres each summer to exploit its vast krill stocks^13,14^.

The relative remoteness of the Southern Ocean meant that humans had limited impacts on the ecosystem until recent centuries^15,16^. However, the discovery of the richness of marine resources in the region resulted in an expansion of hunting in the early to mid-1800s, driving the eradication of many seal rookeries within decades of discovery and leading to the near extirpation of baleen whales by the mid- to late 20th century^17–19^.

The single most important prey item for marine mammals in the Southern Ocean is Antarctic krill (*Euphausia superba*, hereafter krill) and following the massive reduction in baleen whales it was suggested their populations might dramatically increase and drive corresponding increases in other marine predators (the so-called krill-surplus hypothesis^20^). Although there is some support for the krill-surplus hypothesis^21,22^, most research shows that krill populations have declined across the Southern Hemisphere in recent decades^23^. While much of this change has been attributed to climatic changes^24^, there is also growing evidence that the reduction in whale-driven fertilisation of the upper ocean has an important role^13^. Baleen whales recycle iron through consumption and defecation^25^, returning highly labile iron to the surface ocean^26^, crucial for primary productivity in the Southern Ocean where phytoplankton growth is limited by low surface iron concentrations^27^. Estimates suggest that the hunting of baleen whales in the Southern Ocean decreased the recycled iron by an order of magnitude^13^, driving substantial changes in marine ecosystems.

Overlaid on this legacy of exploitation is a rapidly changing physical environment. The Southern Ocean climate is dominated by the Southern Annular Mode (SAM) climatic oscillation, which human-driven stratospheric ozone depletion and rising greenhouse gases have pushed to its most positive sustained state in over a millennium^28,29^. The El Niño–Southern Oscillation (ENSO) is another important climatic driver, and its amplitude has increased since 1960 under greenhouse forcing^30^, with demonstrated impacts on Antarctic seas^31,32^. Together these climate systems change marine communities by altering sea ice extent and the timing of seasonal melting and freezing^33^. These sea ice properties govern the timing and productivity of phytoplankton blooms^34^, drive krill populations across the seasons^24^, determine the light regime reaching benthic macroalgal communities^35^, and impact marine predator populations that rely on sea ice for breeding and foraging^36^. Circumpolar ice extent showed several years of increases following 2010 before shifting into a persistently low state, with a record minimum in 2023 linked to anthropogenic ocean warming^36^.

Environmental changes are particularly pronounced along the Western Antarctic Peninsula, where sea ice responds strongly to combined SAM and ENSO forcing^33^, and the present-day ice season has shortened by more than three months compared to the period 1979–2011^37^. Phytoplankton communities have responded with increases in biomass, bloom duration^38^, and a shift from diatoms towards cryptophytes^39,40^. This shift is important as krill selectively forage on diatoms over cryptophytes^41^ which, in combination with changes to the physical environment, has contributed to a southward shift of krill from the South Western Atlantic towards the Scotia Sea and Antarctic Peninsula^24,42^. The region’s abundant krill also sustained large baleen whale populations, making it one of the most intensively hunted grounds in the Southern Ocean with industrial whaling established at Deception Island in the South Shetlands by the early 1900s^43^. This means that whaling had already depleted predator populations long before ecological surveys documented climate-driven shifts in krill. Thus, climate and whaling effects are superimposed on all observational records, making it difficult to attribute the effect of each stressor with current data that covers only a subset of the impact period. We are therefore unable to untangle the effects of these different anthropogenic impacts and cannot contextualize contemporary biodiversity change against a true historic baseline.

It is now possible to reconstruct past ecosystems beyond the reach of existing observational data by extracting and sequencing environmental DNA (eDNA) from marine sediment cores. Living communities continuously shed DNA into marine sediments, where it can persist for thousands or even millions of years^44,45^. As eDNA is deposited by all life, sampling this archive allows us to reliably reconstruct past community composition over millennia. Ancient eDNA has been used to reconstruct biodiversity changes due to anthropogenic^46^ and natural drivers^47,48^, extending biodiversity records to periods for which no other information exists, and enabling us to measure and assess how communities have changed over time.

Here, we sequenced ancient eDNA from five sediment cores spanning over two centuries to reconstruct whole-community biodiversity across the Western Antarctic Peninsula, comparing these records to historical climate and modelled whale biomass data. We found distinct impacts of anthropogenic climate change and whaling, and revealed an association of whale biomass with water-column feeding animals, consistent with whales sustaining pelagic productivity.

## Results

### Stable communities give way to increasing change after ca. 1900

Across five ^210^Pb-dated cores spanning ca. 1763-2019 we found that western Antarctic Peninsula biodiversity changed little in the oldest parts of our records but shifted rapidly after ca 1900. Change continued through recent decades, producing assemblages increasingly distinct from all earlier states (Fig. 1).

**Figure 1.**
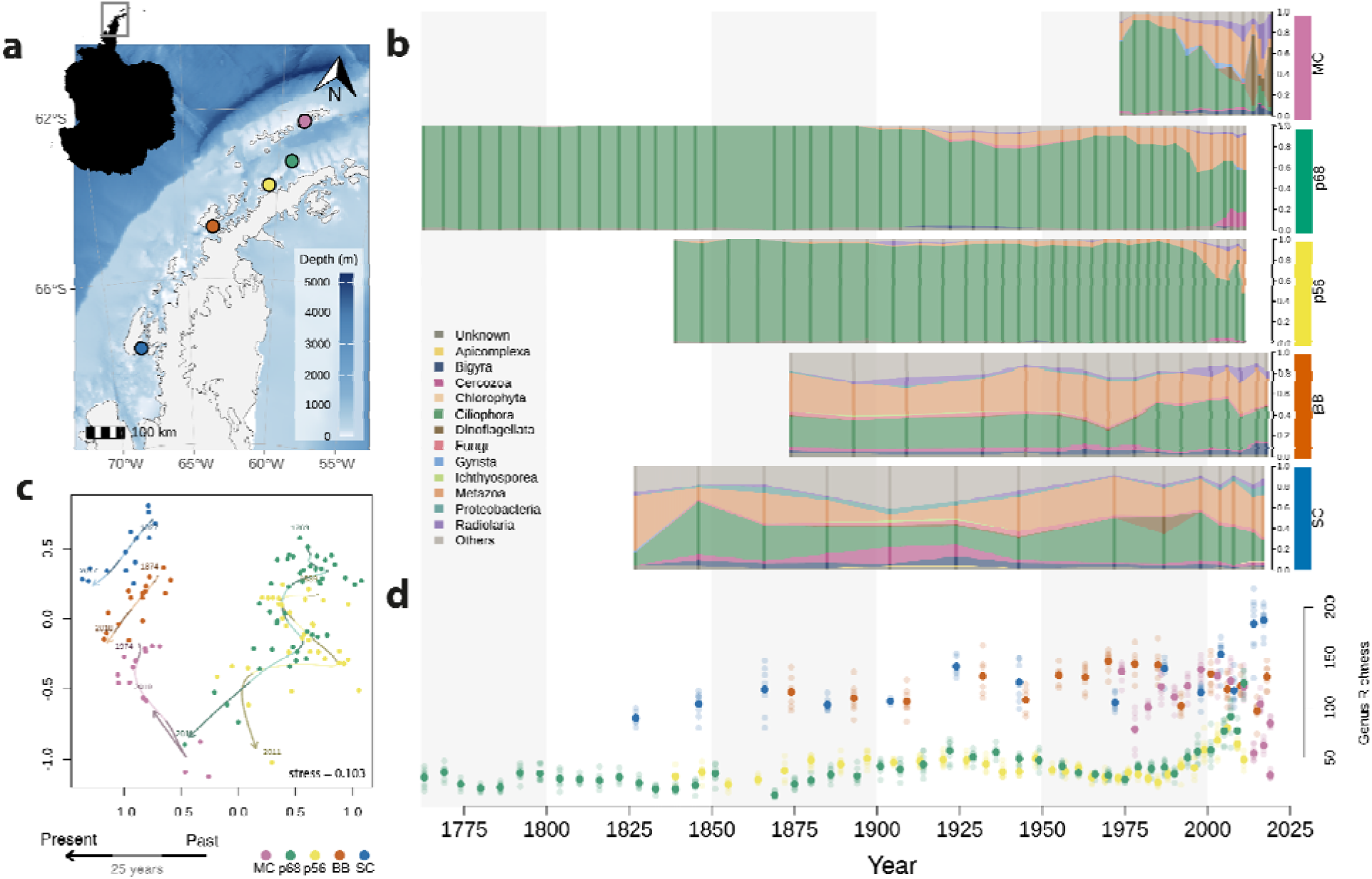
Biodiversity changes from metabarcoding of sediment cores for sites across the Western Antarctic Peninsula. **a** Map of sampling sites shown by coloured points. Bounds shown in the inset silhouette of Antarctica. **b** Relative read abundance of major taxonomic groups (18S PR2^49^ subdivisions), solid bars show sequenced samples. Sites are ordered from top to bottom by latitude (MC - Marian Cove, p68 - Orleans Trough, p56 - Trinity Island, BB - Börgen Bay, SC - Sheldon Cove). Colours on the right axis are consistent across the figure. **c** Community composition shown through non-metric multidimensional scaling ordination based on Jaccard dissimilarities. Lines show a smoothed spline through all data points with dark and light line shades indicating increments of 25 years, the calendar year of the first and last sample is shown above the relevant datapoint. **d** Number of genera (richness) per sample coloured by site, with light-coloured points indicating individual PCR replicates around the mean value shown in dark colours.

We characterised biodiversity through metabarcoding and DNA sequencing of 124 sediment samples amplified at three DNA regions targeting eukaryotes (18S), mammals (mitochondrial 16S) and vertebrates (mitochondrial 12S). Sequencing produced 382.3 million read pairs, with each of the eight PCR replicates generated per sample retaining a mean (± s.d.) of 216,803.5 ± 190,690.6 (18S), 49,114.5 ± 80,723.3 (16S), 27,837.1 ± 54,742.1 (12S) merged sequences after quality control steps (Supplementary Table 1). These sequences generated a total of 17,044 (18S), 283 (16S) and 157 (12S) amplicon sequence variants (ASVs) across the tree-of-life. The 18S region generated ASVs with taxonomic assignments across 53 phyla, 116 classes, 247 families, and 385 genera. The combined 16S and 12S ASVs, which target overlapping taxa, had taxonomic assignments across 25 families and 35 genera.

The 18S DNA relative abundance data (Fig. 1b) showed comparative stability in taxonomic diversity prior to the 1900s. From 1900 onwards, shifts were observed in the northernmost three sites, principally driven by decreases in the protistan clade *Gyrista* (consisting almost entirely of diatom sequences, Supplementary Figure 1). This was followed by even more substantial decreases in *Gyrista* reads in recent decades across sites. Cluster analyses (CONISS) were consistent with these findings: all whole-community zone boundaries (6 of 6) fell after 1900.

Splitting the observations into taxonomic subsets (metazoa, diatom, dinoflagellate and flagellate) revealed the same pattern in community-zone boundaries (Supplementary Figures 1-5). Alpha diversity metrics also revealed substantial change in recent decades (Fig. 1d, Supplementary Figure 6). At every site, either the highest (sites SC, p56 and p68) or lowest (sites MC and BB) genus richness occurred in the last 25 years. Finally, beta diversity showed continuous accelerating compositional change across all sites and dissimilarity metrics (Fig. 1c, Supplementary Figure 7), with ecosystems becoming more compositionally distinct from their historical state through time.

### Quantifying the impacts of anthropogenic drivers

Contemporaneous with our observations, instrumental and modelled climatic records that we compiled showed an increasingly positive Southern Annular Mode (SAM) index^29^ and increased El Niño–Southern Oscillation (ENSO) variability^30^ (Supplementary Figure 8a,c). Average yearly sea ice concentrations^50^ extracted for the sampling sites revealed both declines and increases over the same period (Supplementary Figure 8d), with decreasing yearly sea ice over recent decades in the southernmost site (SC in Fig. 1a) and more open water sites (p56 & p68 in Fig. 1a). Alongside these shifts in the abiotic conditions, modelled baleen whale biomass data^19^ extracted for hunted Southern Ocean species showed a 47-fold reduction (1.55 Mt in 1979 vs. 73.27 Mt pre-1900) across the 20th Century (Supplementary Figure 8e). To evaluate the relative influence of these concurrent pressures on the region’s communities, we further analysed 18S metabarcoding data from the 87 samples (of 124 total) with complete environmental records. We then quantified the effects of SAM, ENSO, sea ice concentration and estimated whale biomass in the Southern Ocean (hereafter whale biomass) on within-sample (alpha) diversity, community composition (beta diversity), and the responses of individual taxa.

Neither climate nor whaling consistently altered the number or evenness of taxa within samples (Hill numbers q = 0-2, Supplementary Figure 9, Supplementary Table 2). Out of all combinations of alpha diversity metrics and drivers tested, three reached nominal significance: SAM was positively associated with richness at both the ASV (q = 0, p = 0.008) and genus level (q = 0, p = 0.025), and whale biomass positively associated with genus-level diversity (q = 1, p = 0.045). ENSO and sea ice concentration did not explain any alpha diversity metrics (all p > 0.05). Local alpha diversity was therefore largely insensitive to both drivers, suggesting that community responses are expressed in which taxa are present, rather than their number or evenness.

Climatic drivers (SAM, ENSO, sea ice concentration) and whale biomass both had a significant joint effect on whole-community beta diversity under abundance-weighted and presence-absence analyses (Bray-Curtis and Jaccard, both p < 0.001; distance-based redundancy analyses conditioned on site, Fig. 2ab, Supplementary Table 3). However, individual drivers were not equally important; SAM and whale biomass were significant under both dissimilarity metrics (p < 0.05), whereas ENSO and sea ice concentration were non-significant throughout (all p > 0.05). Thus, whole-community change was driven principally by an increasing SAM index and whale biomass loss over the study period. Joint species distribution models fitted with Hierarchical Modelling of Species Communities (HMSC^51^), with SAM, ENSO, sea ice concentration and whale biomass as fixed effects and site as a random effect, supported this pattern. Whale biomass had strongly supported associations (posterior support > 0.95) in 38% of metazoan families (21 of 55, Fig. 2c Supplementary Table 4) and 53% of eukaryotic subdivisions (21 of 40, Supplementary Figure 10, Supplementary Table 5) analysed, while SAM responses were more common in subdivisions (40%) than metazoan families (18%). These models also found a significant role for ENSO and sea ice in the responses of some groups (Fig. 2c, Supplementary Table 4-5), not apparent in whole-community analyses. Sea ice concentration notably influenced the majority of metazoan families (58%), with equal numbers of positive and negative responses (Fig. 2c, 16 each direction).

**Figure 2.**
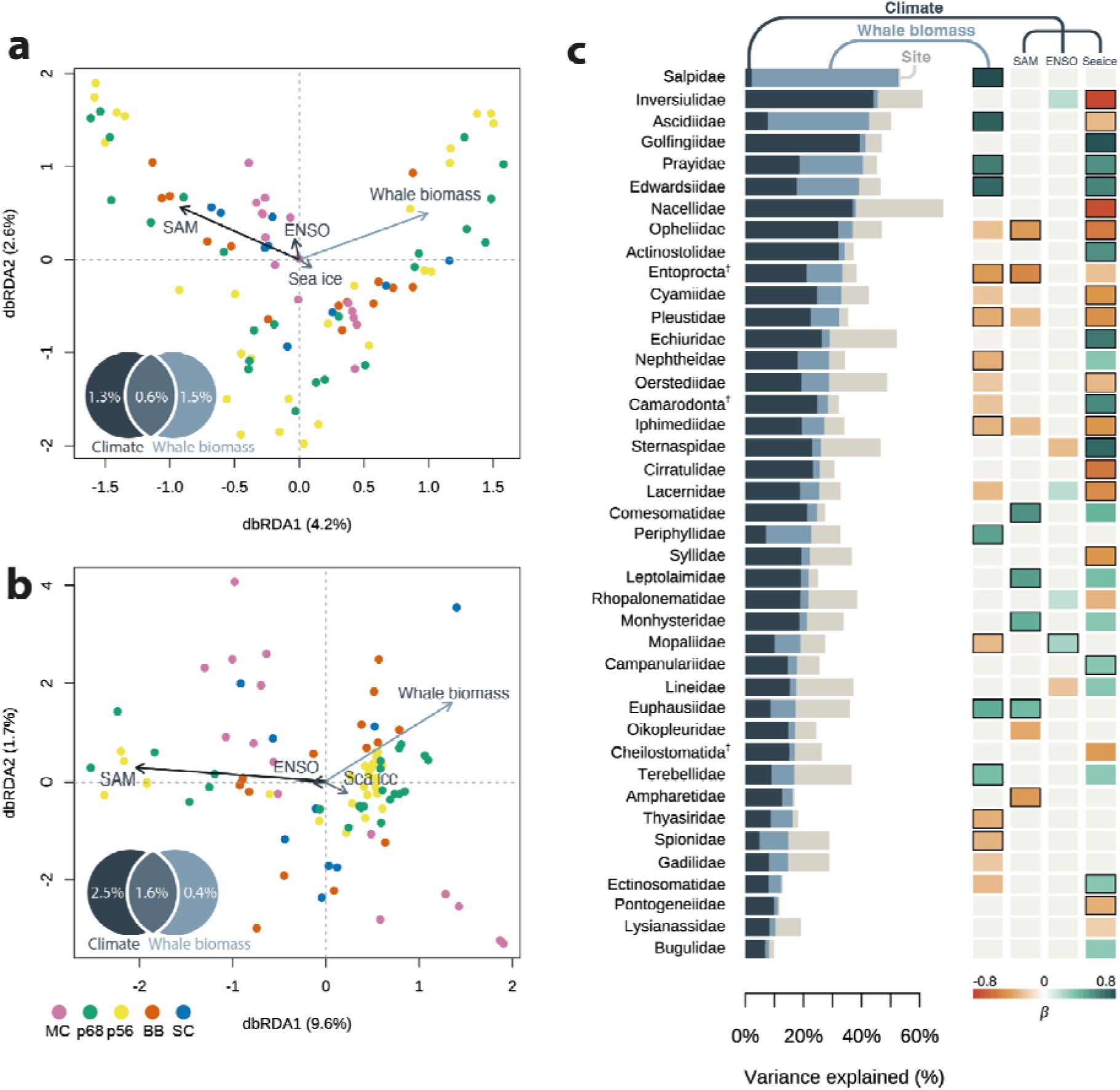
Environmental drivers of community composition across sites. Distance-based redundancy analyses of **a** Jaccard and **b** Bray-Curtis dissimilarities from 18S metabarcoding constrained by whale biomass, the Southern Annular Mode (SAM), El Niño–Southern Oscillation (ENSO) and sea ice concentration, with site included as a conditioning variable. Points represent individual samples, coloured by site. Arrows show the constrained predictors scaled for display; arrow direction indicates the gradient along which each variable structures community variation, and arrow length is proportional to the strength of its association with the ordination axes. Venn diagrams show adjusted R² for each fraction partitioned between whale biomass and the combined climate variables (SAM, ENSO, sea ice). **c** Joint species distribution model (HMSC) fitted to centred log-ratio (CLR)-transformed proportional abundances of metazoan families, with climatic drivers and whale biomass as fixed effects and site as a random effect, families are ordered by the summed explanatory variance of fixed effects. Dagger-labelled taxa indicate taxonomic reassignment to level above family with the lowest rank with Southern Ocean records. Horizontal stacked bars (left) show the proportion of variance in each family attributable to climate, whale biomass, and the site random effect, with the x-axis given as a percentage of total explained variance. Heatmap (right) of mean posterior regression coefficients (β) for each predictor, with red indicating negative and green indicating positive associations on the CLR scale; the colour scale is symmetric around zero and bounded by the maximum absolute β across the displayed taxa. Coloured rectangles indicate posterior probability of effect direction > 0.95, black outlines > 0.99.

Both drivers explained modest, statistically separable fractions of an ecosystem dominated by site differences and residual variation. In the distance-based whole-community analyses (variance partitioning with site as a third explanatory table; Fig. 2ab, Supplementary Table 6), climate uniquely explained 2.5% of abundance-weighted community variation against 0.4% for whale biomass, while the drivers explained approximately similar variance under presence-absence (1.3% against 1.5%), with some overlapping variation across models (0.6–1.6%). This pattern was also observed when splitting the data into metazoan (3.7% versus 1.4%; 0.9% versus 1.0%) and protist subsets (2.1% versus 0.3%; 1.4% versus 1.7%). Among-site differences took the largest unique fraction (37.8% and 17.1%) and residuals remained high (49.7% and 76.1%). These values agreed with variance partitioning of the joint species distribution models, which found that drivers and site effects explained a minority of variation for each taxon (metazoan families mean R² = 0.32, eukaryotic subdivisions 0.38; Supplementary Table 4-5). Variance partitioning (metazoan families and subdivisions respectively) attributed 15.4% and 13.5% to climate, 6.5% and 8.0% to whale biomass, and 9.9% and 16.7% to site effects. High residual variation is characteristic of metabarcoding data^52^ and was broadly consistent across both distance-based (49.7 - 76.1%) and HMSC (61.8 - 68.1%) models.

Climate and whaling produced qualitatively different ecological patterns among the 55 modelled (HMSC) metazoan families (Fig. 2c). Climate drove diverse restructuring, with sea ice affecting 32 families spanning ten phyla in both directions (16 positive, 16 negative). Declines were observed among macroalgae-associated taxa, from limpets (Nacellidae, β = −0.73) to amphipods (Pontogeneiidae, −0.30), and increases among soft-sediment infauna such as peanut worms (Golfingiidae, 0.79). The SAM index was consistently positively associated with free-living nematode families (Comesomatidae, Leptolaimidae, Monhysteridae, 0.43–0.54), while ENSO had minor effects compared to other drivers. The whale biomass response, in contrast, separated animals according to feeding traits (Fig. 3). The six largest positive coefficients were all animals that capture food from the water column: salps (Salpidae, β = 0.80), ascidians (Ascidiidae, 0.74), burrowing anemones (Edwardsiidae, 0.70), siphonophores (Prayidae, 0.62), scyphozoan jellyfish (Periphyllidae, 0.48) and krill (Euphausiidae, 0.45), while negatively associated families were predominantly small benthic invertebrates. Across all modelled families, water-column feeders responded more positively to whale biomass than other taxa (Wilcoxon p = 0.031) and the average difference in whaling coefficients between the two groups was 0.21 (95% CI 0.14–0.29, Supplementary Table 7). No climate variable showed comparable response structure (SAM, ENSO and sea ice coefficient differences all between −0.03 and 0.01, 95% CIs spanning zero, all Wilcoxon p > 0.05). A presence-absence metazoan family model agreed throughout: no family differed in the direction of its whaling association between models (11 of 24 strongly supported in both), and water-column feeders again responded more positively than other taxa (Wilcoxon p = 0.044), with a mean difference in whaling beta coefficients of 0.17 (95% CI 0.05–0.29, Supplementary Figure 11-12, Supplementary Table 8).

**Figure 3.**
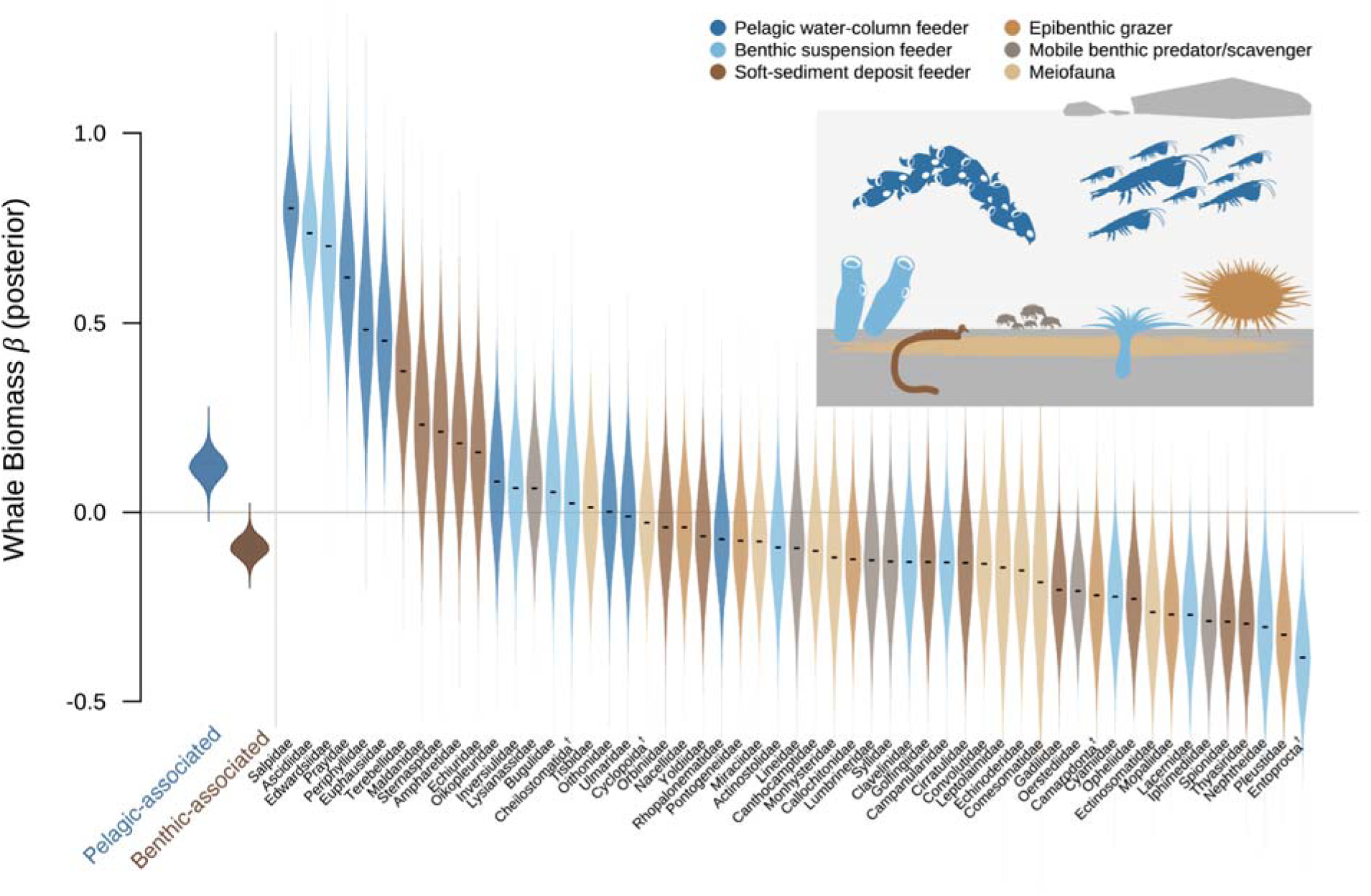
Whale biomass positively associates with metazoans which feed from the water column. Joint species distribution modelling effect sizes (beta posterior) of whale biomass on metazoan family centred log-ratio (CLR) transformed read abundances from 18S metabarcoding. Violin plots show the spread of beta posteriors over the MCMC estimates for each family, with the mean shown with a black line and the colour indicating feeding strategy as shown in the top right schematic. The distribution of mean beta posteriors across all MCMC estimates for pelagic and benthic associated feeders is shown on the left. Dagger-labelled taxa indicate taxonomic reassignment to level above family with the lowest rank with Southern Ocean records.

Eukaryotic subdivisions supported the water-column to sediment division in responses (Supplementary Figure 10). Whale biomass positive associations were broadly planktic (cryptophytes, β = 0.64; radiolarians, 0.64; haptophytes, 0.37) and declines were sediment-associated heterotrophs (cercozoan flagellates, −0.49), while the largest SAM responses were benthic amoebae (Discosea, 0.69) and foraminifera (0.48). The Gyrista response ran counter to expectations under whale-mediated iron fertilisation, with a negative association with whale biomass (−0.24) as well as with SAM (−0.33).

### Changes across the food web from plankton to whales

The impacts of human drivers can be seen in changing detection patterns of blue whales (*Balaenoptera musculus,* combined 16S and 12S data), krill (*Euphausiidae*, 18S data) and phytoplankton (18S data) in Figure 4. Blue whales (Fig. 4a) showed declining eDNA detections across the time series, with the highest detection frequency in cores sampled from deeper sites (p56 & p68) prior to the 1900s, and a substantial decrease in occupancy across all sites in the 20th century. Detections correlated with modelled biomass estimates (Supplementary Figure 13). Krill showed mixed patterns over time between sites, with decreases in the deep-water sites (p68 and p56) spanning the first half of the 1900s, followed by sharp increases from ca. 1975 onwards. The most southerly site (SC) showed an increase from ca. 1940 onwards, and the most northerly site (MC) a decrease from the start of the 2000s, agreeing with observations of krill populations moving southwards in response to anthropogenic climate change^42^. Finally, mirroring the observed decrease in diatoms in the taxonomic overview (Fig. 1b), we observed a sharp decrease in the ratio of sequences from diatoms to flagellate phytoplankton from ca. 1975 onwards in the three most northerly sites (MC, p68, p56).

**Figure 4.**
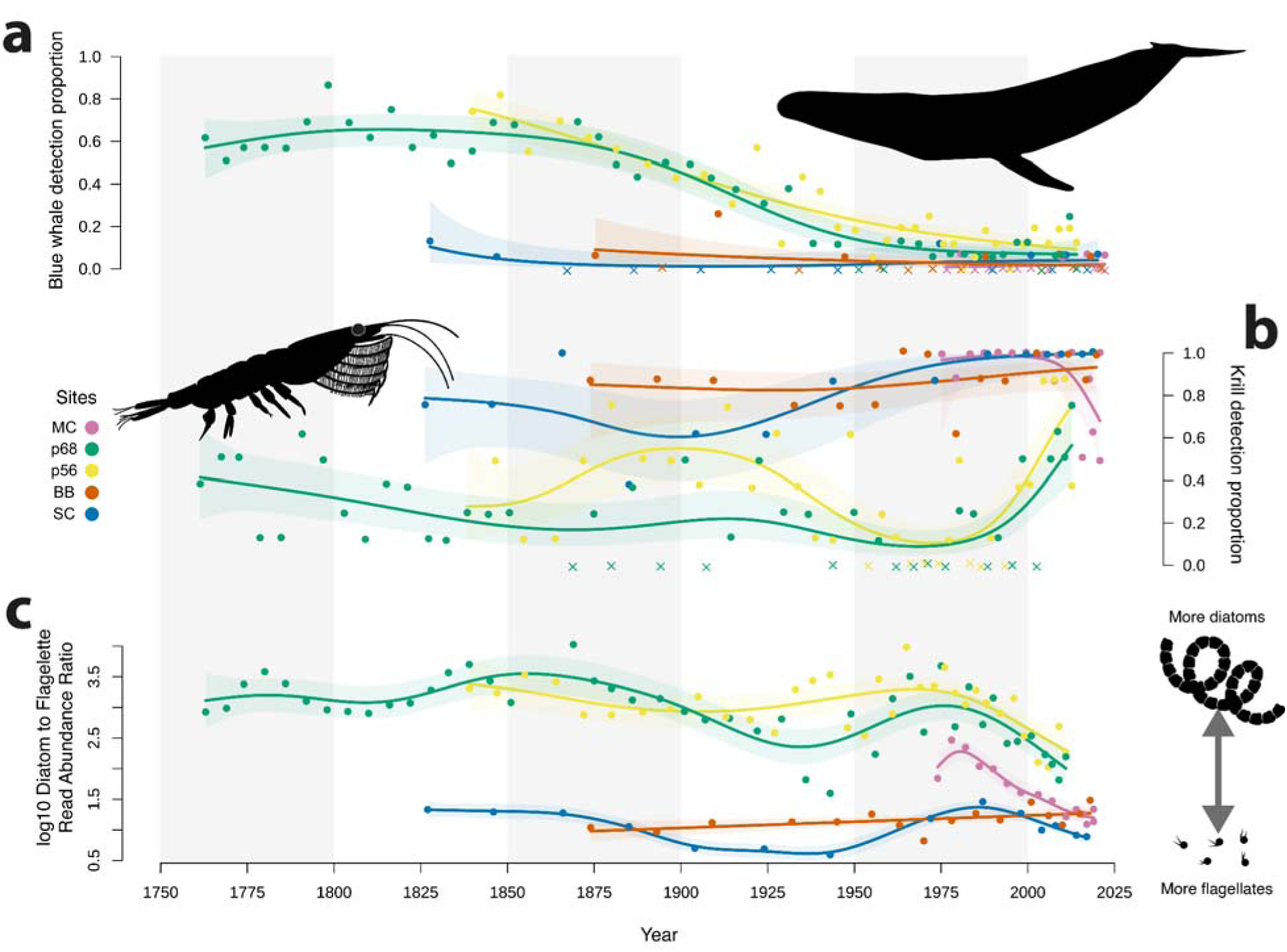
Changes in selected Antarctic marine taxa over time. Lines are generalised additive model (GAM) fits for each site, and shaded bands are 95% confidence intervals, point and line colours indicate sites. **a** Blue whale (Balaenoptera musculus) detection frequency across 16S and 12S metabarcoding PCR replicates. **b** Krill (family Euphausiidae) detection frequency across 18S metabarcoding PCR replicates. **c** Ratio of diatom to flagellate relative read abundance (log10). Diatoms are the classes Bacillariophyceae, Mediophyceae and Coscinodiscophyceae. Flagellates are plastid-bearing taxa with a flagellate life stage. Mixed groups (e.g. Dinophyceae, Chrysophyceae), aplastidic taxa and non-marine classes are excluded. Supplementary Figure 14 shows the same ratio with the flagellate group widened to all plastid-bearing marine non-diatom taxa.

## Discussion

Our ancient eDNA time series revealed a Southern Ocean ecosystem under substantial anthropogenically forced biodiversity change across 200 years. We found a key role for both climate change and historic whaling in driving responses at both the whole-community level and for individual taxa. Moreover, the community change attributed to different drivers was taxon-specific, with diverse turnover across the tree-of-life in response to climatic drivers, and a response suggestive of a loss of pelagic productivity associated with whaling. These results provide both a glimpse into pre-impact marine Antarctic ecosystems and a rich picture of how anthropogenic forces have driven biodiversity towards novel modern states.

Our understanding of biodiversity change in the Southern Ocean principally comes from biomonitoring and satellite records spanning recent decades^38,40,53^, all of which commenced after the onset of industrial whaling^17^. With eDNA-based reconstructions we extended these observations, confirming pervasive climate-linked change but showing that climate shifts and whaling restructured communities in different ways. We observed a decline in blue whale detections in our cores mirroring declines reconstructed across the whole Southern Ocean^19^ (Supplementary Figure 13a). However, we did not find as strong a pattern for fin whales (*Balaenoptera physalus*, Supplementary Figure 13b) whose pre-impact and human-extracted biomass is estimated to have been at least as large as that of blue whales^19^. This highlights an important limitation of eDNA metabarcoding data: some taxa are detected poorly or not at all. We thus focus on those taxa which were detected, acknowledging that our inferences are drawn from a subsample of the whole community.

Whaling had a significant impact on marine biological communities across our time series, but we additionally identify that this response is associated with taxon feeding traits. As whale biomass fell from near-unexploited levels, communities shifted away from both pelagic (salps, siphonophores, krill) and benthic (ascidians, burrowing anemones) water-column feeding organisms towards diverse mixed benthic assemblages, including polychaete worms and molluscs. As the balance between suspension and deposit feeders on the Antarctic shelf tracks the supply of surface-derived food^54^, this is the pattern expected if whale loss reduces pelagic productivity. A link between whale abundance and pelagic productivity is central to the hypothesis that whales act as ocean fertilisers^13,25^. Until now, support has come from enrichment experiments^55^, the high lability of iron in whale faeces^26^, and ecosystem modelling^13,56^. Our time series data add a new line of evidence that whales shape the wider ecosystem through fertilisation.

Krill were positively associated with whale biomass, consistent with the hypothesis that whale-mediated nutrient recycling supported larger krill stocks prior to exploitation^25^, and with reported declines since the whaling period^24^. Recent work has sought to understand observations of shifting dominance of the two major pelagic grazers, salps and krill, in the Southern Ocean^57^. Our data reveal a positive association of both groups with whale biomass suggesting that recent shifts towards a salp dominated ecosystem^23^ may be pre-dated by declines in all pelagic grazers due to whaling. This association supports modelling results that suggest a pre-whaling Southern Ocean with substantially larger carbon export compared to the present day^58^.

Antarctic diatoms are a key link in recycling iron back into the food web; however, our models (HMSC, Supplementary Figure 10) showed a negative relationship between whale biomass and Gyrista sequence abundance (principally diatom sequences, Supplementary Figure 1). Given the overall abundance and sharp recent decrease of diatoms over our time series (Fig. 1b), the HMSC identifies an unexpected influence when the effects of other drivers are accounted for. As the dominant group, Gyrista’s relative abundance is most sensitive to increases in other taxa. Its sharpest declines also occurred after ca. 1975, when whale biomass was slightly recovering and SAM was most positive (Supplementary Figure 8). The negative coefficient may therefore reflect an interplay between the read dominance of this taxon and the timing of these trends, rather than evidence against whale-mediated iron fertilisation. Artificial ocean iron enrichment experiments have found that dissolved inorganic iron increases primary productivity, particularly favouring diatoms in the Southern Ocean^59^. However, we have only limited experimental data confirming that phytoplankton incorporate biogenic iron from whale faeces^55^ and no work has yet performed enrichment experiments with iron supply with whale faecal-like properties. The Western Antarctic Peninsula shelf is also one of the more iron-replete regions of the Southern Ocean^60^, so other whale-derived nutrients may be important for plankton fertilisation, as seen at lower latitudes^61^.

Climatic shifts drive ecosystem change across a large number of variables, for example, temperature, salinity, turbidity and ice scour, each of which impact different taxa positively or negatively^36,62^. In our data this is observed in the large spread of responses across phyla, with climate driving a large proportion of the total change (Fig. 2ab), and diverse taxa responding to changes in the physical environment (Fig. 2c). Increasing krill detections at the southernmost site and decreasing detections at the northernmost site in recent decades (Fig. 4b) support the documented recent poleward contraction of krill along the Western Antarctic Peninsula^24,42^. We also observe a sharp shift in diatom to flagellate phytoplankton sequences in the three northernmost sites across the last 50 years (Fig. 4c), in line with observations of phytoplankton community shifts away from diatom dominated assemblages^39,40^. Moreover, as we find this shift to be unique to recent decades, it shows that climate-driven phytoplankton changes are without precedent across at least two hundred years.

Crucially, our data cover over two hundred years, offering a hitherto unseen baseline of Antarctic biodiversity. While we reveal a marine ecosystem under persistent change (Fig. 1), we are unable to assess if the observed ecosystem states would fall within the possible range for the environment without human impacts. Moreover, our evidence does suggest that recent decades are particularly divergent from observed historical states, but the sequence of impacts (whaling followed by climate change) makes it difficult to rule out an interaction between stressors which may influence our ability to understand the unique importance of each driver^63^.

Here we have focused on the influence of climate change and whaling, but it is feasible that other important drivers may be unaccounted for in our models, biasing our ability to uniquely attribute impact to stressors. An important example is sealing: we detected five seal species in our data (16S and 12S metabarcoding, Supplementary Figure 15) and observed a recent increase in the Antarctic fur seal (*Arctocephalus gazella*) suggestive of recovery. However, as we do not observe any declines correlating with major sealing in the early 1800s^43^, we did not attempt to attribute biodiversity change to sealing, and thus cannot rule out impact in our data. Another key anthropogenic change in recent decades is the krill fishery established during the 1970s. This fishery has operated well below (<10%) the precautionary catch limit for the Southwest Atlantic^24^, which is a fraction of the total standing stock, but we cannot rule out local effects on our krill signal as much of the fishery is concentrated around the Antarctic Peninsula. While the fishery and our interpretation of krill patterns are focused on *Euphausia superba*, an important limitation of our metabarcoding data is our lack of intrafamily resolution to identify krill taxa within *Euphausiidae*. Thus, although *Euphausia superba* likely represents the bulk of our signal, other krill species may influence the observed patterns.

Future Antarctic seas will host larger whale populations as their numbers recover, but these animals will encounter the increasing effects of human-driven climate change^64^. While our data suggest greater whale biomass will support biodiversity associated with enhanced primary productivity, it remains unclear how this will interact with the continued impacts of climate change. Despite the substantial ecosystem changes documented here, Antarctica remains one of the regions least impacted by humans today ^65^. Its conservation should therefore be a priority, and ancient eDNA records, like those presented here, offer a long-term baseline to contextualise recovery and future change.

## Methods

### Sediment records

Five sediment records from across the Western Antarctic Peninsula were selected to represent broad-scale ecosystem changes. Three multicores were collected in 2019 during the JR19002 cruise on the RRS *James Clark Ross*^66^; SC0 from Sheldon Cove, Rothera, Adelaide Island (364 m depth, −67.5986 - 68.1015); BB0 from Börgen Bay, Anvers Island (313 m depth, −64.7645 - 63.4713); and MC0 from Marian Cove, King George Island (225 m depth, −62.2199 - 58.8172). These records are all from coastal sites with no benthic scour from nearby glacial outflow and are referred to in this work as SC, BB and MC respectively. These cores were extruded onboard at 0.5 cm intervals for the first centimetre from the surface, 1 cm intervals for samples from 1 cm to 4 cm, and 2 cm for the remainder of the core. Subsequent analyses were performed on these slices. Extruded slices were stored in aluminium foil lined petri dishes at −20 °C until further processing. Two multicores were collected in 2016 during the PS97 (ANT-XXXI/3) cruise on the RV *Polarstern*; PS97/056-1 from the east of Trinity Island (633 m water depth, −63.7570 - 60.4418); and PS97/068-2 from the Orleans Trough (1992 m water depth, −63.1675 - 59.3020), referred to throughout as p56 and p68 respectively. These cores were frozen whole at −20 °C until later processing.

### Chronology

Age models for cores from each cruise were independently constructed, with full details in Mason et al.^67^ and Vorrath et al.^68^ for the JR19002 and PS97 cruises respectively, with analytical details summarised below.

For cores SC, BB and MC, ^210^Pb and ^214^Pb activities were determined by gamma spectroscopy on sealed, equilibrated samples counted for a minimum of 48 hours at the University of Plymouth Consolidated Radioisotope Facility. Excess ^210^Pb (^210^Pb_ex) was calculated as the difference between total ^210^Pb and supported ^210^Pb (inferred from ^214^Pb). Age-depth models and mass accumulation rates were calculated using the Constant Rate of Supply (CRS) models^69,70^. For further details see Mason et al.^67^.

For cores p56 and p68, ^210^Pb_ex activities were determined as the difference between total ²¹ Pb and ^226^Ra. Activities were measured by alpha spectroscopy for p56 and gamma spectroscopy for p68. Ages were calculated using the CRS model, with full analytical details provided in Vorrath et al.^68^.

For all cores the mean sedimentation rate of the oldest three samples was used to extrapolate the age-depth model below the measured values; these dates are thus provisional extrapolations and should be interpreted as such.

### Metabarcoding

All laboratory steps prior to PCR were conducted in specialised ancient DNA facilities according to standard protocols for ancient sediments^71^. The whole frozen cores were sliced while frozen and subsampling of all material was conducted as follows. After core slices reached room temperature, material was subsampled using sterile single-use tools only from the centre of the slice, ensuring no contaminating material on the outside of the slice was sampled. These subsamples were frozen at −20°C until DNA extraction. A total of 124 samples were extracted from the five cores: SC (n = 14), BB (n = 16), MC (n = 15), p56 (n = 34) and p68 (n = 45). For each sample 0.4 g of material was processed using the MagAttract PowerSoil Pro kit (Qiagen N.V., Netherlands) (protocol version date 09/2022) with the following modifications. QSB1 binding buffer was modified following Holman et al.^47^ by combining QSB1 buffer, 5 M sodium acetate and 5 M sodium chloride at a ratio of 189:10:1, respectively. To increase the retrieval of smaller DNA fragments the post-digestion lysate was combined with modified QSB1 binding buffer and MagAttract Suspension G beads at a 1 to 3 volume ratio, processing 240 μL of lysate for purification. This purification was performed in duplicate to produce two DNA extracts for each sample. The final elution was performed using 60 μL of solution C6. An extraction negative control was included for each run of 16 samples and included in all subsequent steps.

Prior to metabarcoding all samples were tested for inhibition by spiking each sample into a well characterised probe-based qPCR reaction, ensuring less than 1 cycle threshold difference between the unspiked and spiked reactions to ensure no inhibition. Reaction details are provided in Holman et al.^47^. Three DNA regions were used to characterise past biodiversity with metabarcoding. A variable length (90-140 bp excluding primers) portion (V9) of the eukaryotic nuclear 18S ribosomal RNA gene (Euk1391f: 5’ GTACACACCGCCCGTC 3’ ^72^, EukBr: 5’ TGATCCTTCTGCAGGTTCACCTAC 3’ ^73^). A ca. 97 bp portion of the mitochondrial 12S ribosomal RNA gene (12S-V5_F: 5’ TAGAACAGGCTCCTCTAG 3’, 12S-V5_R: 5’ TTAGATACCCCACTATGC 3’ ^74^) and a ca. 95 bp portion of the mitochondrial 16S ribosomal RNA gene (16Smam1 5’ CGGTTGGGGTGACCTCGGA 3’, 16Smam2 5’ GCTGTTATCCCTAGGGTAACT 3’ ^75^). These regions are referred to throughout as 18S, 12S and 16S, respectively. Forward and reverse primers incorporated unique nucleotide tags of 8 base pairs in length (>3 differences between tags) on the 5’-end. Optimal PCR cycle numbers were determined by qPCR as detailed in Holman et al.^47^. For each sample, eight independent PCR replicates were conducted in 20 µL reactions consisting of 1 × AmpliTaq Gold 360 Master Mix, 0.5 μM forward and reverse primer, and 2 μL eDNA template. PCR was conducted with an initial denaturation at 95°C for 10 minutes followed by cycles (26 for 18S, 40 for 12S and 16S) of 95°C for 30 s, variable annealing temp (57°C for 18S, 51°C for 12S, 59°C for 16S) for 30 s, and 72°C for 1 min. Negative extraction controls and negative PCR controls (2–5 negative controls for every 40 PCR replicates) were amplified alongside experimental samples and included in sequencing. Following PCR amplification, products were visualized on a 2% agarose gel and equimolarly pooled according to the strength of the target DNA band (15 μL for very light/no band, 10 μL for medium strength band and 5 μL for strong band), and 15 μL of PCR product from extraction negative controls and PCR negative controls were added. Negative control sample read summaries are shown in Supplementary Table 1b.

A total of 12 amplicon pools per metabarcoding primer set were purified with the MagBio HighPrep PCR clean-up system (MagBio Genomics, Inc, Gaithersburg, MD, USA) using the manufacturer-provided protocol with a 1:1 (18S) or 0.8:1 (12S and 16S) bead to amplicon pool ratio. The purified amplicon pools were built into Illumina sequencing libraries using the TagSteady library build protocol^76^, which minimises tag-jumping. Unique dual indexes (different index for each forward and reverse adaptor, each index used only once) of 10 nucleotides in length were used for indexing pools and the libraries were sequenced across 2 SP flowcell lanes (250 bp paired-end) with 10% PhiX library spike-in on the Illumina NovaSeq 6000 Instrument.

### Bioinformatics

Illumina demultiplexed amplicon pool libraries were demultiplexed into PCR replicates using Cutadapt (v.5.0)^77^ by matching the forward and reverse primers (including tags) with one incorrect base permitted. This matching was repeated in the opposite orientation with the same stringency, as the library build method creates libraries in both orientations. Sequences in different orientations were then processed independently using DADA2 (v.1.35.8)^78^ in R (v.4.5.0)^79^ using the default settings unless detailed below. As each sequencing run has a unique error profile the following steps were conducted for each run independently. Demultiplexed reads were filtered using the filterAndTrim function with the following parameters ‘minLen = 50, maxN = 0, maxEE = c(1,1), truncQ = 2’. The NovaSeq instrument bins per base-pair sequence quality data, resulting in poor error model fitting using the learnErrors function, therefore an error model enforcing monotonicity was used as outlined in the supplied bioinformatic script. Following amplicon sequence variant (ASV) generation, the resultant data tables from each demultiplexing orientation were combined, summing ASVs with 100% identity (once oriented in the reverse complement). Sequencing runs were then combined into a single dataset, identical ASVs were merged and chimeras removed. Singleton observations and any ASV occurring in only a single replicate were removed. ASVs outside of the expected marker length (18S 90-200 bp, 16S 89-120 bp, 12S 89-110 bp) were removed. Any observation from the experimental samples with fewer reads than the maximum number of reads per ASV across all negative controls was set to zero.

ASVs were taxonomically assigned using blastn (v.2.16.0+)^80^ against the NCBI nt database (downloaded 15 Feb 2024) to return 200 hits (-num_alignments 200) per ASV; these were then parsed using a custom R script that uses a lowest common ancestor approach to assign a taxonomy to each ASV (ParseTaxonomy, doi:10.5281/zenodo.4671710) with high quality assignments reported in the results above 95% coverage and 99% sequence similarity. The assignTaxonomy function from DADA2 with the protist-oriented PR2 database (v.5.1.0)^49^ was used to generate a second set of assignments as this database uses a different taxonomic classification system and provides high-confidence higher level taxonomic assignments (e.g. protist/metazoa). All species level assignments (e.g. marine mammals) were manually confirmed using the online NCBI blastn search tool (last accessed July 2025) by ensuring each assignment differed by at least 1bp to congeners.

### Environmental data

We characterised Southern Annular Mode (SAM) variability up to the present by combining the Marshall Index observational data^81^ and multi-proxy reconstruction (Abram et al.^28^). Over the 1957–2007 overlap, the observations and model outputs differed by a constant offset, which we subtracted to place the reconstruction onto the scale of the observational data.

We characterised El Niño–Southern Oscillation (ENSO) variability using monthly Niño 3.4 sea surface temperature anomalies (5°N–5°S, 170–120°W) from the NOAA Extended Reconstructed SST (version 5)^82^. Data were summarised as a November-January mean assigned to the calendar year of January capturing the boreal event peak.

Estimated sea ice coverage for each site was extracted from sea ice reconstructions^50^ by finding the nearest available datapoint for each site within seasons and averaging across the year to give an estimated yearly percentage. The location used in each case is shown in Supplementary Figure 16.

Estimates of whale abundance were extracted from Greenspoon et al.^19^, which combines ecosystem models and catch data to estimate total biomass across the Southern Ocean. Taxa subject to whaling and found in both the modelled and metabarcoding datasets were extracted for analysis. The final dataset included blue (*Balaenoptera musculus*), fin (*Balaenoptera physalus*), humpback (*Megaptera novaeangliae*) and sei (*Balaenoptera borealis*). The earliest data point for each species represents the first documented whaling catch, which occurred at different periods for each species. Models including whale biomass as a covariate used the summed biomass of all four whales as a proxy for all whaling effort, with the first value for each species extended backwards to cover the period 1775 to 2020. Given the absence of fine scale whaling records across the study time period, and the proximity of the sampling region to major whaling grounds^17,20,43^, these Southern Ocean values represent a proxy for local whaling effort. All biodiversity analyses formally assessing the role of SAM, ENSO, sea ice and whale biomass were conducted on a data subset with entries for all environmental parameters, parameters were scaled and rolling 3-year mean values for SAM and ENSO used for analyses.

### Ecological analyses

Taxonomic diversity barplots were generated using relative proportion normalised datasets with taxa corresponding to less than 1% of data in each sample collapsed into ‘other’ for visualisation. Compositional transitions within sites were identified by stratigraphically constrained hierarchical clustering (CONISS, function *chclust* in rioja v1.0-7^83^) on Bray–Curtis dissimilarities of age-ordered samples, with the number of significant zones determined by with a broken-stick model. This was applied separately to the full dataset and to metazoan, diatom, dinoflagellate and flagellate (excluding dinoflagellates) families. Detection proportion was calculated across all technical replicates and is presented as a proxy for DNA concentration in plots above following Baños et al.^84^.

For alpha diversity estimates, PCR replicates with fewer than 30,000 reads were discarded and the remaining PCR replicates rarefied to the minimum remaining read count using *rrarefy* in R (vegan *v.2.6.10*). Hill numbers of order 0 (richness), 1 (exponential of Shannon entropy) and 2 (inverse Simpson) were calculated and the mean over 100 rarefaction draws taken as the estimate. Alpha diversity was calculated for ASVs and also for genus richness by summing ASVs with identical taxonomic assignments (>60 bootstrap uncertainty cutoff, unassigned ASVs discarded). Linear mixed models (lme4 v.1.1.37)^85^ were fitted to each Hill number with SAM, ENSO, site sea ice concentration and summed whale biomass as fixed effects (∼ SAM + ENSO + SeaIce + whale) and coring site as a random intercept.

Non-metric multidimensional scaling ordinations were constructed using both Jaccard and Bray-Curtis dissimilarities on relative proportion transformed datasets using *metaMDS* in vegan with t*rymax* set to 200. Distance-based redundancy analyses were conducted using *dbrda* in R (vegan), with separate models fitted for Jaccard and Bray-Curtis dissimilarities from relative proportion transformed ASV tables. The effect of site was partialled out using *Condition()* and marginal significance was assessed by permutation (10 permutations, blocked by site). The comparative effect of climatic variables (SAM, ENSO and sea ice) and whale biomass was further estimated using *varpart* (vegan) under the default parameters. Beta-diversity analyses were repeated on metazoan and protist subsets (Supplementary Tables 3 & 6).

To further assess the response of individual taxonomic groups to climate variability and whaling we used joint species distribution models fitted with Hierarchical Modelling of Species Communities (*Hmsc in R v.3.3.7*) ^51,86^. For eukaryotic subdivisions, ASVs with PR2 subdivision bootstrap ≥ 60 were collapsed to subdivision (53 after filtering) with only eukaryotic subdivisions (40 total) presented in the results. Prokaryotic subdivisions recovered through off-target amplification were retained in all models, as they contribute to the compositional transformation and joint model structure, but are not interpreted individually given the limited and biased recovery of prokaryotes by the 18S marker. For metazoan families, ASVs assigned to Metazoa by PR2 with family-level bootstrap of 100 and carrying an NCBI family name were collapsed to family (names follow the NCBI assignment); families present in fewer than 10% of samples were discarded, retaining 55. For abundance-based analyses, ASV counts were normalised to relative proportions across the full eukaryotic community before collapsing to the focal taxonomic level, zeros were replaced with 0.65 times the minimum non-zero value^87^ and data were centred-log-ratio (CLR) transformed prior to modelling with a normal distribution. For the presence-absence model, a family was scored present where any constituent ASV was detected; families present in all samples were additionally excluded as invariant, retaining 54, and modelling used a probit distribution. All models included the four scaled drivers as fixed effects (∼ SAM + ENSO + sea ice + whale biomass) and site as a random effect. Models were fitted using six MCMC chains with a transient of 100,000 iterations and 4,000 retained samples per chain at a thinning rate of 100. Convergence was assessed via effective sample size and Gelman-Rubin diagnostics on the Beta, Omega and Sigma parameters (Supplementary Table 9). Effects were considered strongly supported where the posterior probability of the coefficient’s sign exceeded 0.95. Explanatory power was quantified per taxon as R² (Tjur R² and AUC for the probit model; evaluateModelFit), and variance components are reported as proportions of total variance, computed per taxon as the partitioned fraction (computeVariancePartitioning; climate = SAM, ENSO and sea ice) multiplied by that taxon’s R² and averaged across taxa.

All 55 modelled metazoan families were screened against the Register of Antarctic Marine Species^88^ using the REST API accessed on 25 Aug 2026. Families without Southern Ocean records were relabelled at the nearest higher rank with records, unrecognised families were traced to the taxonomically accepted family. Families were assigned to six broad ecological groups reflecting habitat and dominant feeding mode. To test whether responses were structured by feeding ecology, we calculated the difference in mean coefficient between water-column-feeding families and all other families for each posterior sample. We included all modelled families, regardless of whether their individual responses were strongly supported. This gave a posterior distribution for the difference, which we summarised as its mean and 95% credible interval. Wilcoxon rank-sum tests on posterior mean coefficients were additionally calculated as a frequentist alternative.

## Data & Code Availability

All processed data and code are available at doi.org/10.5281/zenodo.23808604 for review, and a post-review version will be archived prior to publication. Raw sequence data will be uploaded at the European Nucleotide Archive prior to publication.

## Acknowledgements

We extend our thanks to the Globe Molecular Biology Labs and the GeoGenetics Sequencing Core at the University of Copenhagen for their essential work and support in laboratory work and DNA sequencing. We thank Bent Petersen for his continued support with the Mjölnir high-performance computing cluster. We thank Christian Carøe for discussions on early versions of this work.

