## Supplementary Information for "Ancient environmental DNA reveals the distinct fingerprints of whaling and climate change on Antarctic marine biodiversity"

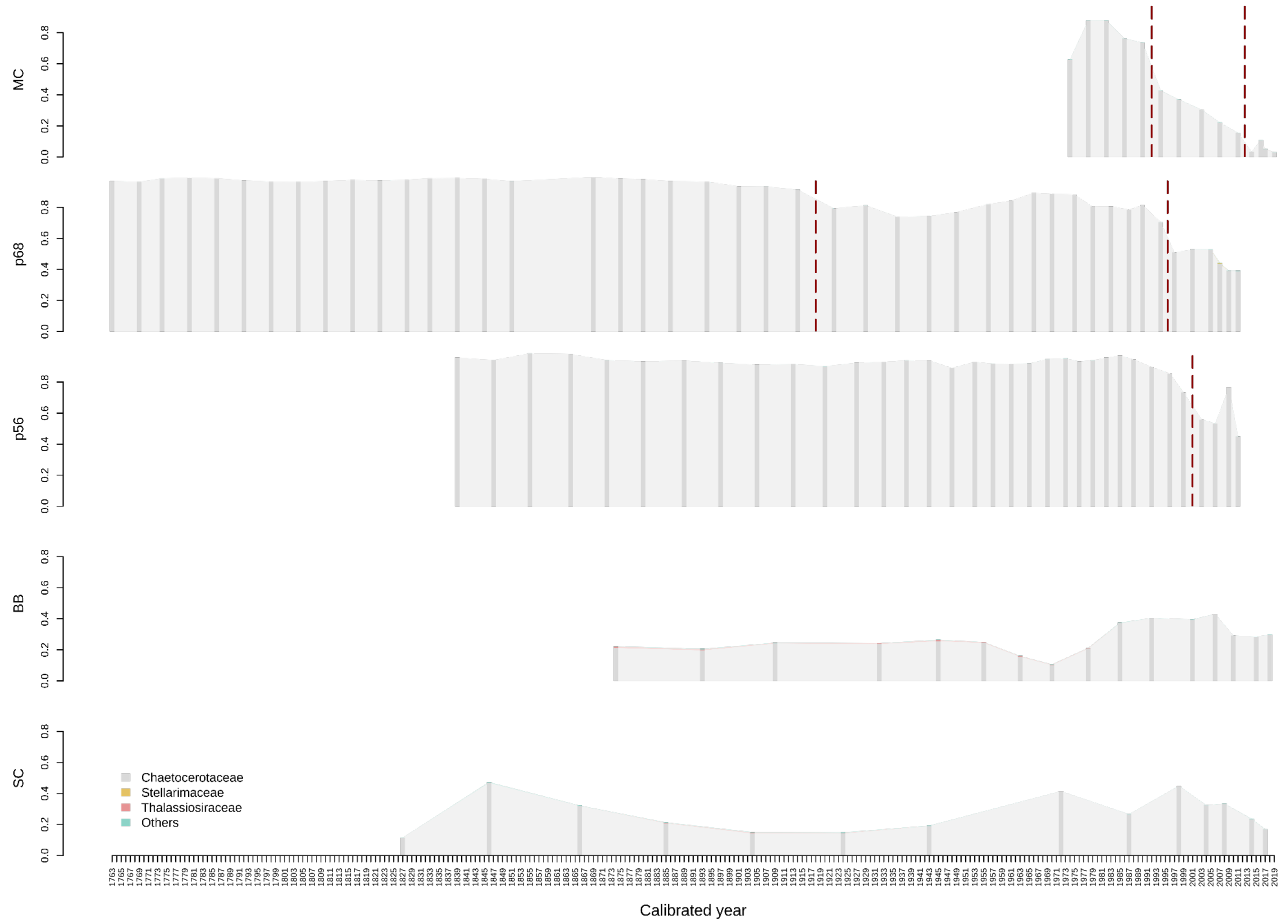


**Supplementary Figure 1:** Relative read abundance of diatom families through time at each of the five coring sites, from 18S rDNA amplicon data. Diatom ASVs (class-level assignment with bootstrap support above 80) were converted to proportions of total eukaryotic reads, aggregated to family (unassigned reads grouped as 'Unknown'), and families reaching at least 1% relative abundance in any sample were retained. Stacked bars give the proportion of total reads, with shaded ribbons connecting adjacent sampled years. Barplots per site are ordered from most northerly to most southerly, matching the layout in the main figures. Dashed red vertical lines mark stratigraphically constrained zone boundaries identified by CONISS cluster analysis on Bray–Curtis dissimilarities of the site's diatom composition.

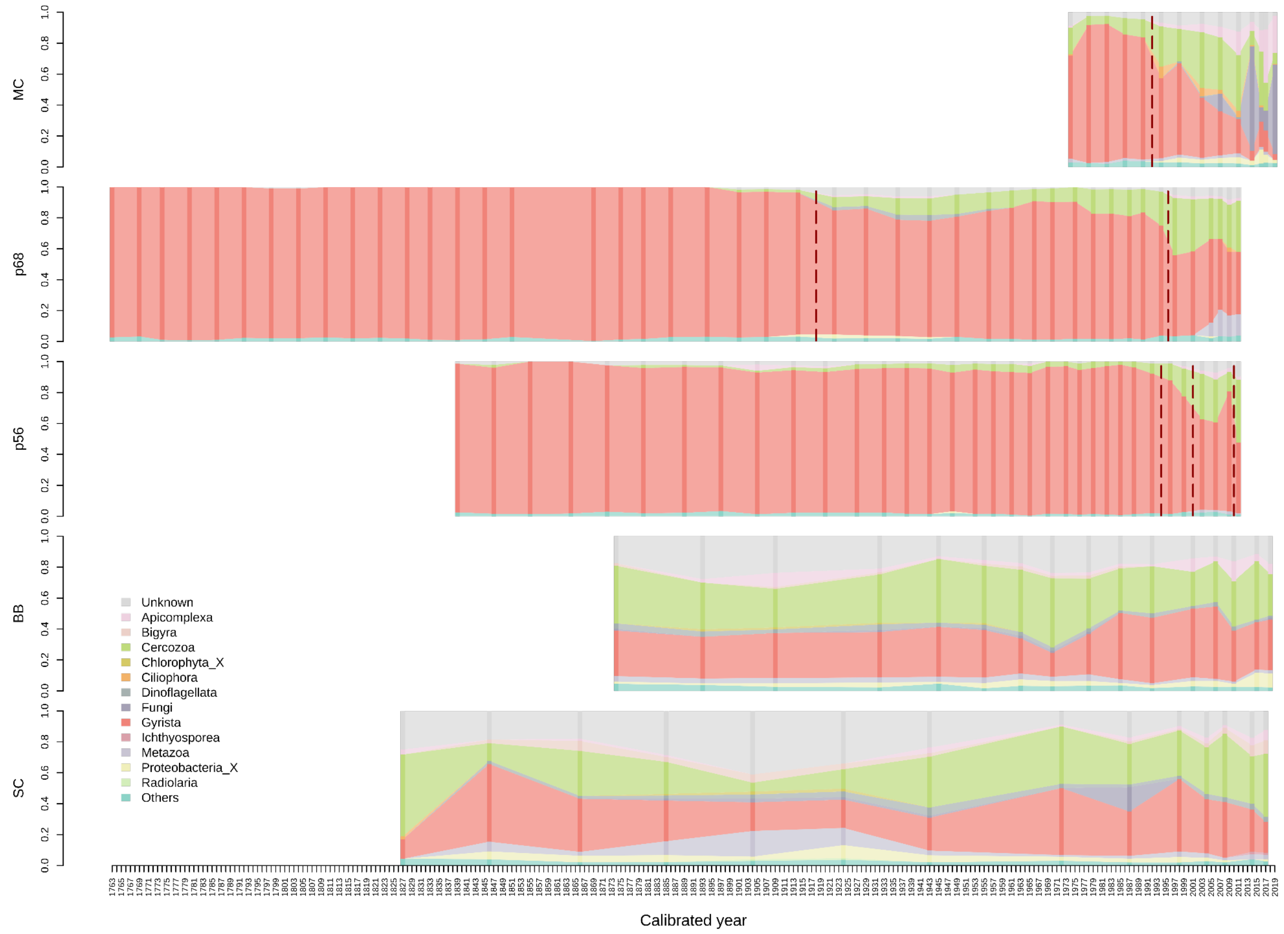


**Supplementary Figure 2:** Relative read abundance of eukaryotic subdivisions through time at each of the five coring sites, from 18S rDNA amplicon data. Reads were converted to proportions and aggregated to PR2 subdivision without a bootstrap support threshold (unassigned reads grouped as 'Unknown'), and subdivisions reaching at least 1% relative abundance in any sample were retained. Stacked bars give the proportion of total reads and therefore sum to one within each sample, with shaded ribbons connecting adjacent sampled years. Barplots per site are ordered from most northerly to most southerly, matching the layout in the main figures. Dashed red vertical lines mark stratigraphically constrained zone boundaries identified by CONISS cluster analysis on Bray–Curtis dissimilarities of the site's whole-community composition.

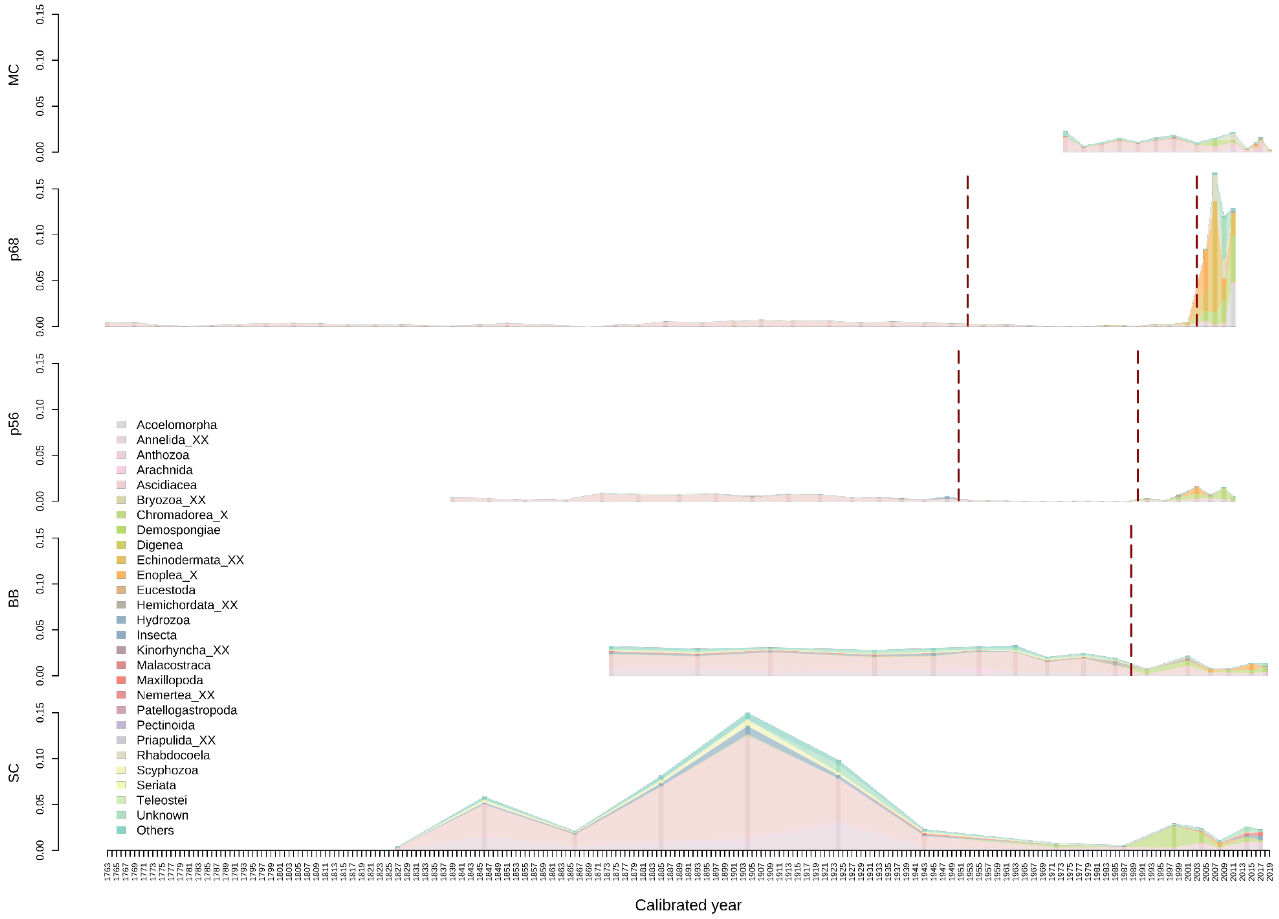


**Supplementary Figure 3:** Relative read abundance of metazoan families through time at each of the five coring sites, from 18S rDNA amplicon data. Metazoan ASVs (subdivision-level assignment with bootstrap support above 80) were converted to proportions of total eukaryotic reads, aggregated to family (unassigned reads grouped as 'Unknown'), and families reaching at least 1% relative abundance in any sample were retained. Stacked bars give the proportion of total reads, with shaded ribbons connecting adjacent sampled years. Barplots per site are ordered from most northerly to most southerly, matching the layout in the main figures. Dashed red vertical lines mark stratigraphically constrained zone boundaries identified by CONISS cluster analysis on Bray–Curtis dissimilarities of the site's metazoan composition.

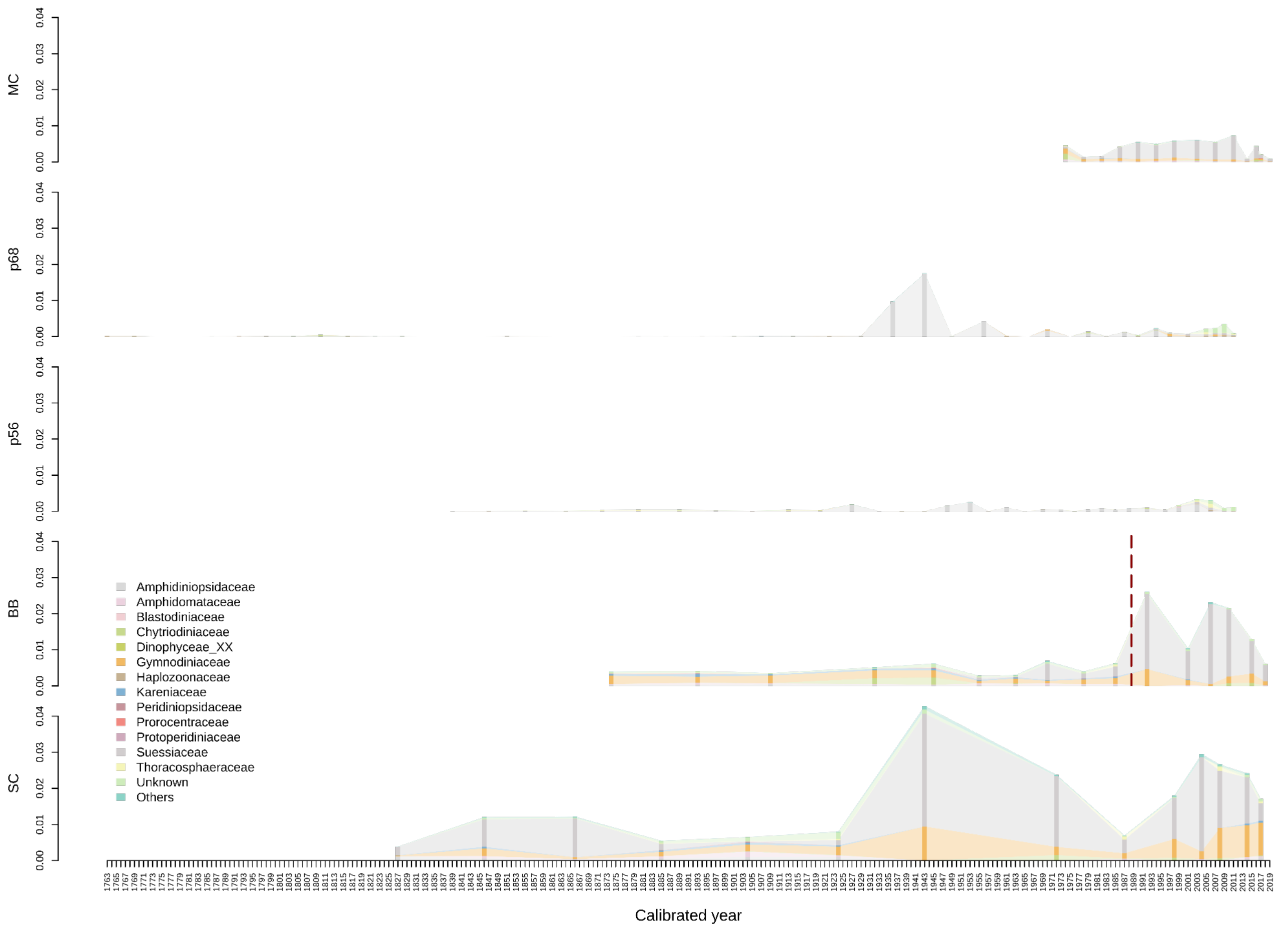


**Supplementary Figure 4:** Relative read abundance of dinoflagellate families through time at each of the five coring sites, from 18S rDNA amplicon data. Dinoflagellate ASVs (class-level assignment with bootstrap support above 80) were converted to proportions of total eukaryotic reads, aggregated to family (unassigned reads grouped as 'Unknown'), and families reaching at least 1% relative abundance in any sample were retained. Stacked bars give the proportion of total reads, with shaded ribbons connecting adjacent sampled years. Barplots per site are ordered from most northerly to most southerly, matching the layout in the main figures. Dashed red vertical lines mark stratigraphically constrained zone boundaries identified by CONISS cluster analysis on Bray–Curtis dissimilarities of the site's dinoflagellate composition.

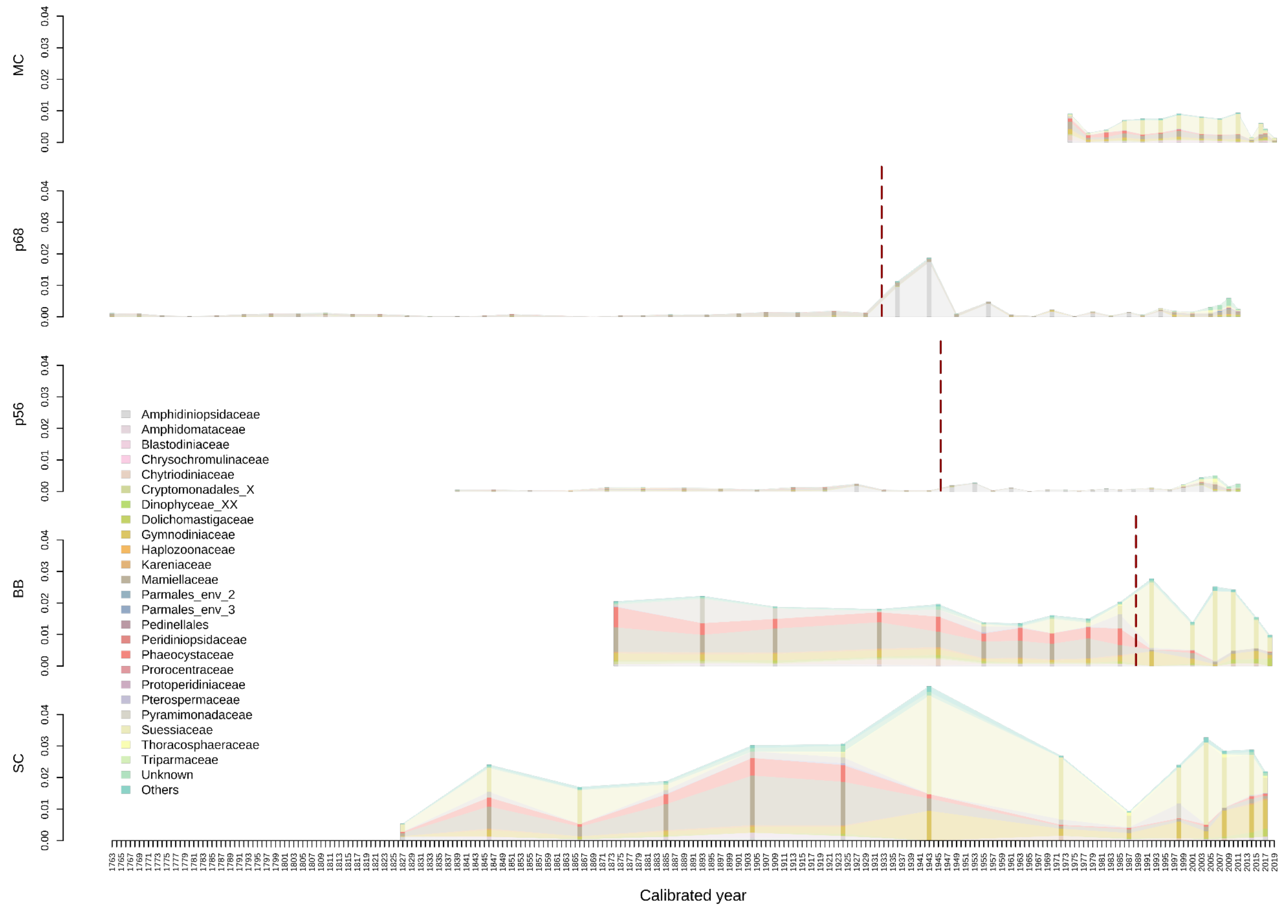


**Supplementary Figure 5:** Relative read abundance of flagellate families through time at each of the five coring sites, from 18S rDNA amplicon data. Flagellate ASVs (class-level assignment with bootstrap support above 80) were converted to proportions of total eukaryotic reads, aggregated to family (unassigned reads grouped as 'Unknown'), and families reaching at least 1% relative abundance in any sample were retained. Stacked bars give the proportion of total reads, with shaded ribbons connecting adjacent sampled years. Barplots per site are ordered from most northerly to most southerly, matching the layout in the main figures. Dashed red vertical lines mark stratigraphically constrained zone boundaries identified by CONISS cluster analysis on Bray–Curtis dissimilarities of the site's flagellate composition.

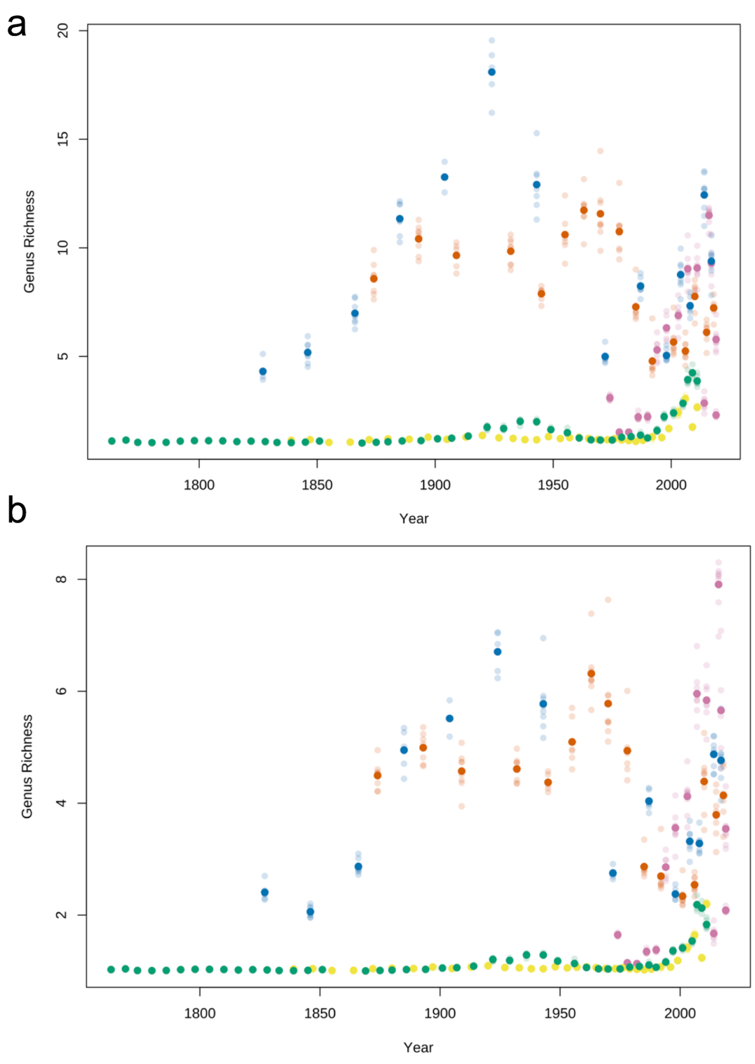


**Supplementary Figure 6:** Within-sample (alpha) diversity through time for each site: Hill numbers of **a** order 1 (exponential Shannon) and **b** 2 (inverse Simpson) computed on rarefied reads at genus level, with points for individual PCR replicates coloured according to site matching main figures.

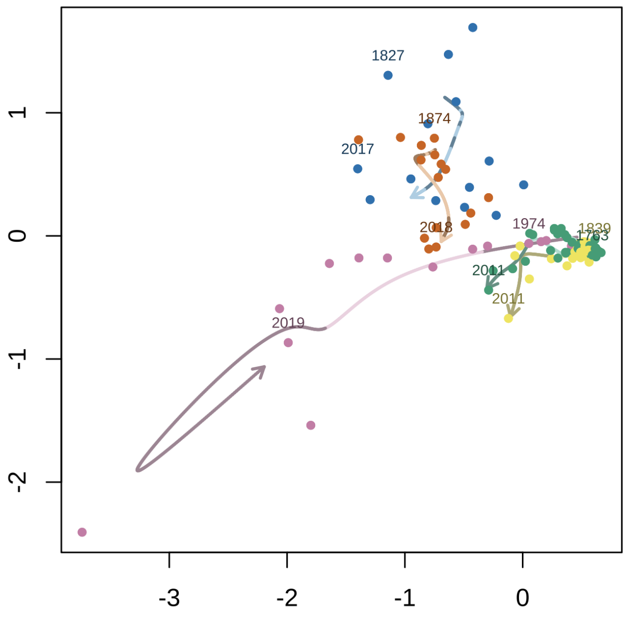


**Supplementary Figure 7:** Community composition shown through non-metric multidimensional scaling ordination on Bray-Curtis dissimilarities for 18S data. Lines show a smoothed spline through all data points with dark and light line shades indicating increments of 25 years, the calendar year of the first and last sample is shown above the relevant datapoint. Ordination stress = 0.082.

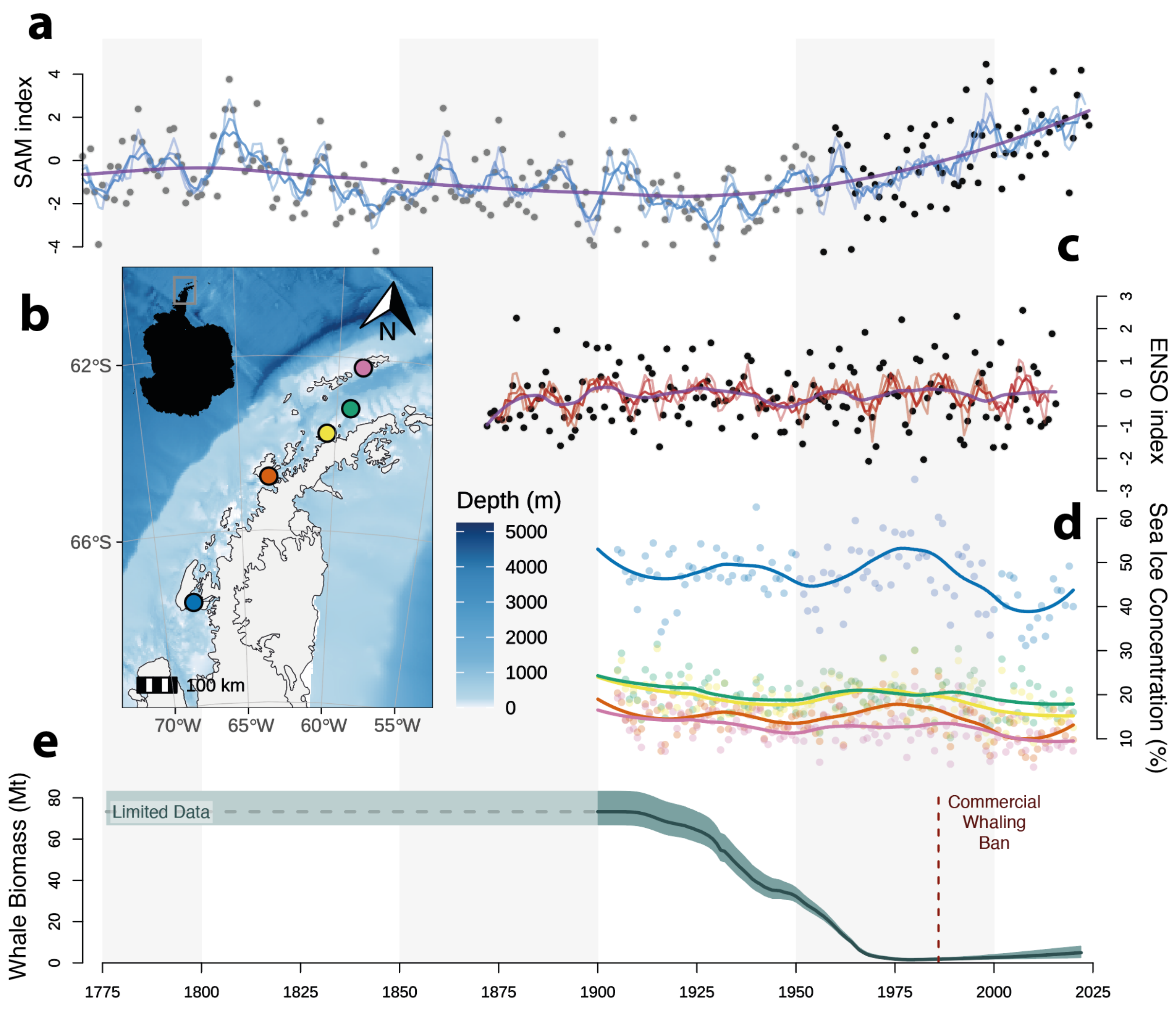


**Supplementary Figure 8: Environmental changes across the Western Antarctic Peninsula**
**a** Yearly mean Southern Annular Mode (SAM) variability from model [1] (grey) and observation [2] (black) data with 3,5, & 7 year rolling averages (blue) and a loess smooth in purple. **b** Map indicating core sampling locations. **c** Yearly mean (Nov-Jan) El Niño-Southern Oscillation (ENSO) variability [3] (black) with 3, 5, & 7 year rolling average (red) and a loess smooth in purple. **d** Average yearly sea ice concentration (%) [4] with loess smooth fitted line, colours match sites in b **e** Modelled summed biomass [5] (millions of metric tonnes) across the Southern Ocean for Blue, Fin, Humpback and Sei whales. Background grey bars indicate 50 year increments.

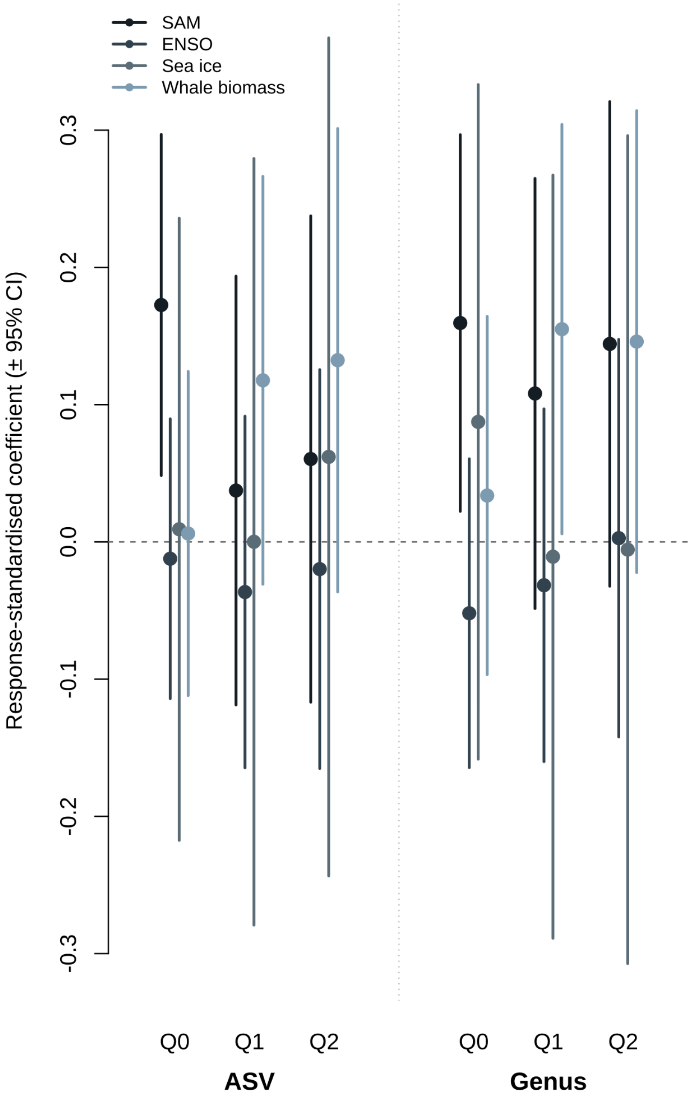


**Supplementary Figure 9:** Fixed-effect coefficients from linear mixed models of alpha diversity against the four scaled drivers. Alpha diversity is measured by Hill numbers of order 0 (richness), 1 (exponential Shannon entropy) and 2 (inverse Simpson concentration), computed on rarefied reads at ASV and genus level and averaged across PCR replicates. Each metric was modelled separately as diversity ~ SAM + ENSO + sea ice + whale biomass + (1 | site), fitted by REML with core site as a random intercept (lme4). All four predictors were centred and scaled to zero mean and unit variance, and coefficients were additionally divided by the standard deviation of their response. Points are point estimates and vertical bars are 95% Wald confidence intervals; colour denotes the driver, the horizontal dashed line marks zero (no effect).

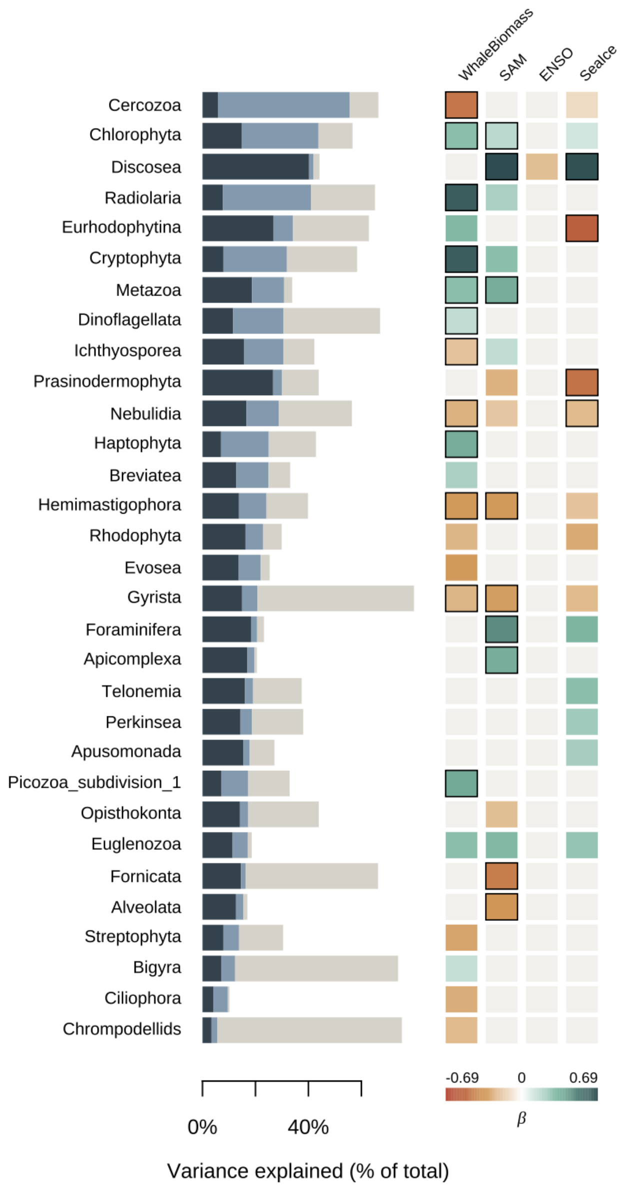


**Supplementary Figure 10:** Joint species distribution model (HMSC) fitted to CLR-transformed proportional abundances of subdivisions from 18S eDNA metabarcoding. Climatic drivers and whale biomass are modelled as fixed effects and site as a random effect. Subdivisions are ordered by the summed variance explained by fixed effects. Taxonomy follows the PR2 database designations. Horizontal stacked bars (left) show the proportion of variance in each subdivision attributable to climate (dark blue), whale biomass (light blue), and the site random effect (grey), with the x-axis given as a percentage of total explained variance. Heatmap (right) of mean posterior regression coefficients (β) for each predictor, with red indicating negative and green indicating positive associations on the CLR scale; the colour scale is symmetric around zero and bounded by the maximum absolute β across the displayed taxa. Coloured rectangles indicate posterior probability of effect direction > 0.95, black outlines > 0.99.

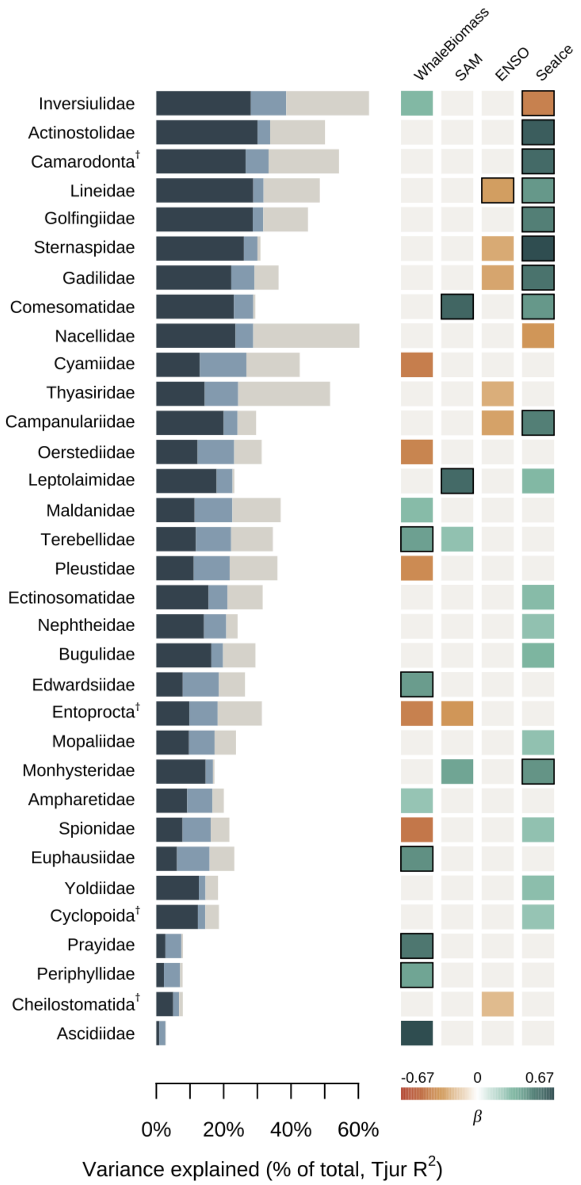


**Supplementary Figure 11:** Joint species distribution model (HMSC) fitted to presence-absence (probit link) metazoan families data from 18S eDNA metabarcoding. Climatic drivers and whale biomass are modelled as fixed effects and site as a random effect. Families are ordered by the summed variance explained by fixed effects. Taxa labelled with '†' indicate taxonomic reassignment to level above family with the lowest rank with Southern Ocean records. Horizontal stacked bars (left) show the proportion of variance in each family attributable to climate (dark blue), whale biomass (light blue), and the site random effect (grey), with the x-axis given as a percentage of total explained variance. Heatmap (right) of mean posterior regression coefficients (β) for each predictor, with red indicating negative and green indicating positive associations on the probit scale; the colour scale is symmetric around zero and bounded by the maximum absolute β across the displayed taxa. Coloured rectangles indicate posterior probability of effect direction > 0.95, black outlines > 0.99.

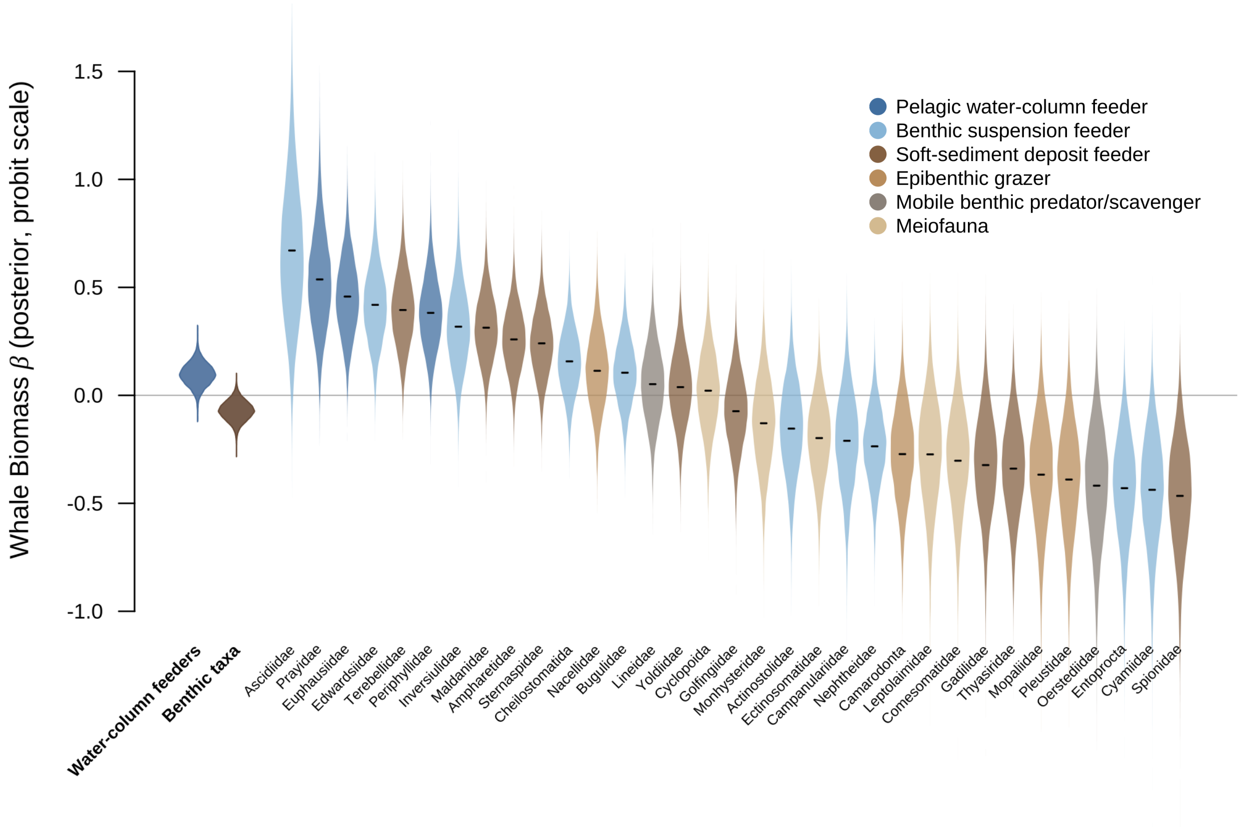


**Supplementary Figure 12:** Joint species distribution modelling (HMSC) effect sizes (beta posterior) of whale biomass on metazoan family presence-absence from 18S metabarcoding. Violin plots show the spread of beta posteriors over the MCMC estimates for each family, with the mean shown with a black line and the colour indicating feeding strategy as shown in the top right schematic. The distribution of mean beta posteriors across all MCMC estimates for pelagic and benthic associated feeders is shown on the left.

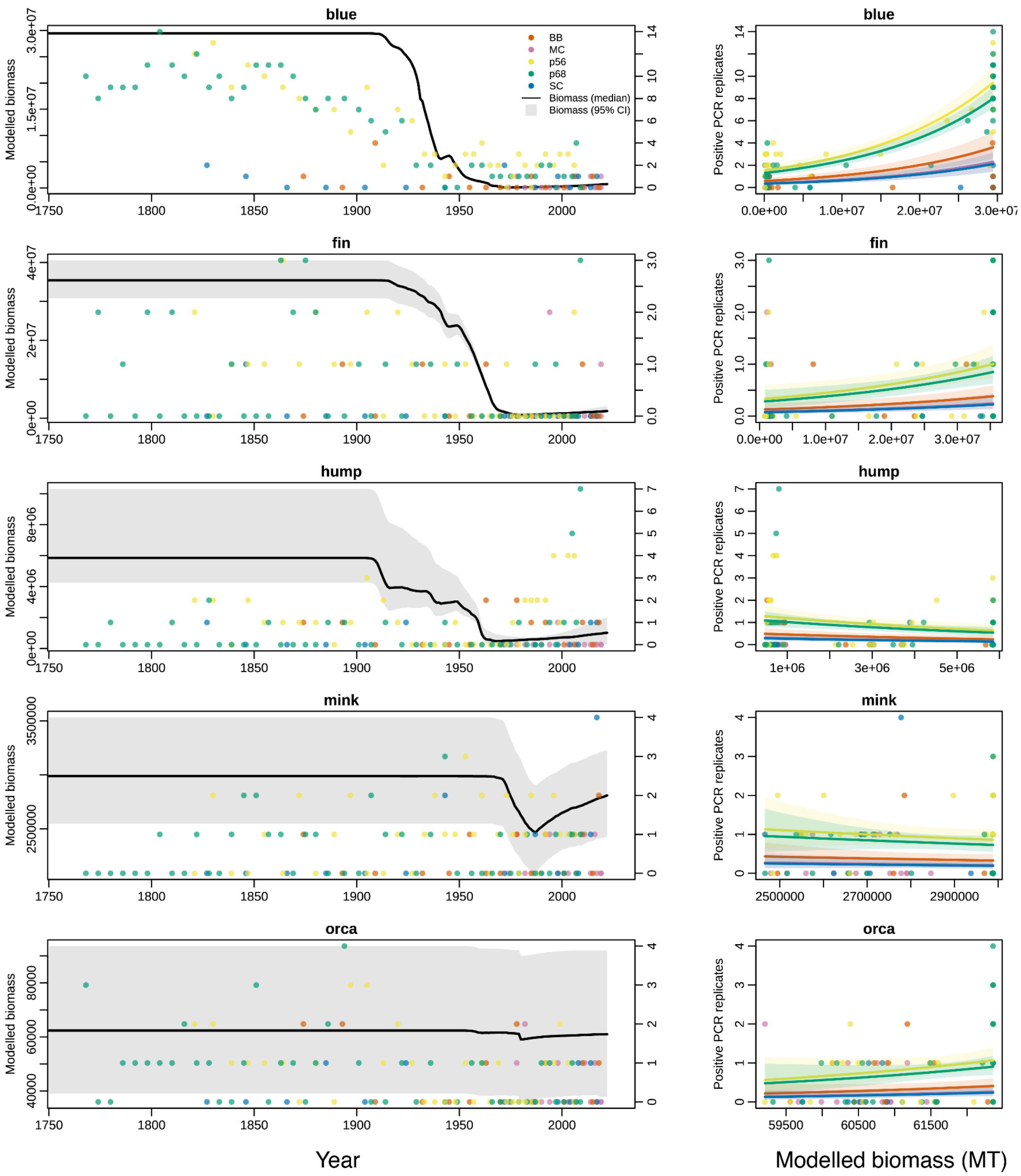


**Supplementary Figure 13:** Modelled whale biomass (in millions of tonnes) and sedimentary eDNA detections for five Southern Hemisphere whale species, by row from top: blue (*Balaenoptera musculus*), fin (*B. physalus*), humpback (*Megaptera novaeangliae*), Antarctic minke (*B. bonaerensis*) and killer whale (*Orcinus orca*). Biomass trajectories are from Greenspoon et al. (2025). Detections are positive PCR replicates summed across mitochondrial 16S and 12S metabarcoding assays from Antarctic marine sediment cores, with colours indicating site. Left: median modelled biomass (black line, 95% CI grey) and positive replicates per dated sample (points, right axis), 1760–2025. Right: positive replicates as a function of modelled biomass, fitted with a negative binomial GLM (log link). The model included species, biomass (standardised within species) and their interaction, plus site as a fixed effect. Lines show predicted means per site, with 95% confidence intervals shaded.

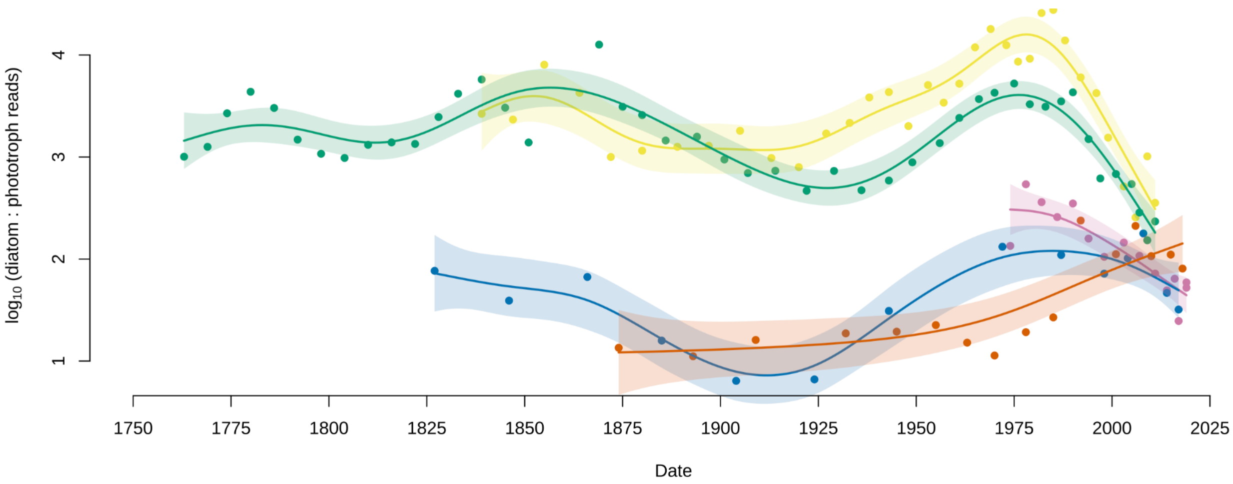


**Supplementary Figure 14:** Diatom to non-diatom phototroph ratio over time, using a broader definition than Fig. 4c. Points are samples, coloured by site as in Fig. 4. Lines are GAM fits for each site (smoothness chosen by REML), and shaded bands are 95% confidence intervals. The ratio is diatom relative read abundance divided by the relative read abundance of all plastid-bearing marine non-diatom taxa, log10 transformed. Unlike Fig. 4c, the non-diatom group includes non-motile taxa, such as coccoid picoplankton (e.g. *Bathycoccus, Ostreococcus, Chloropicophyceae, Pelagophyceae*) and unassigned haptophytes and *mamiellophytes*. Mixed trophic groups (e.g. *Dinophyceae, Chrysophyceae*), aplastidic taxa, uncultured clades of unknown morphology and non-marine classes are excluded, as in Fig. 4c.

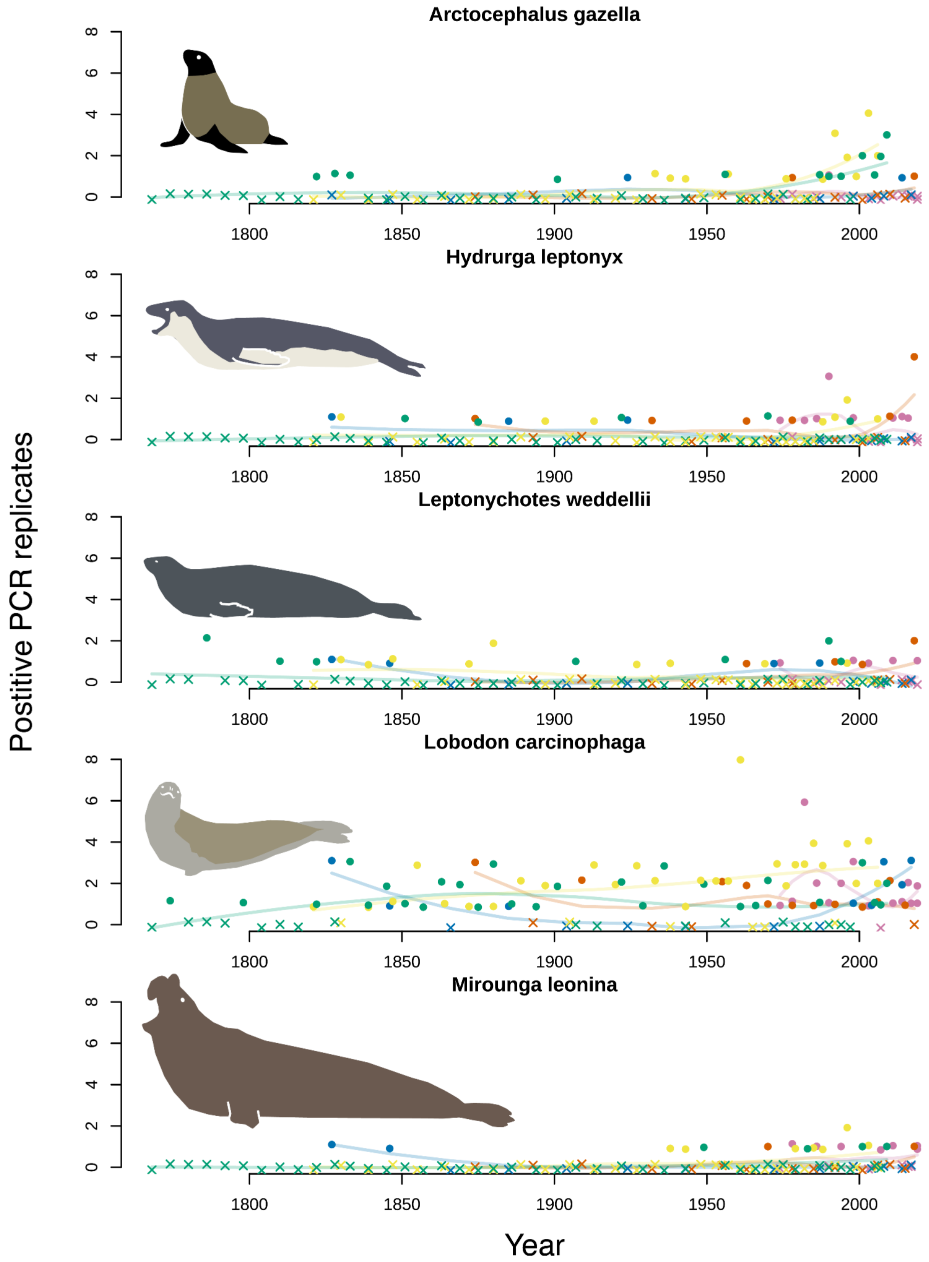


**Supplementary Figure 15:** Detection of seals in eDNA metabarcoding data, pooled across the mammal/vertebrate 16S and 12S markers. Panels show Antarctic fur seal (*Arctocephalus gazella*), leopard seal (*Hydrurga leptonyx*), Weddell seal (*Leptonychotes weddellii*), crabeater seal (*Lobodon carcinophaga*) and southern elephant seal (*Mirounga leonina*). Each point is one sample, showing the number of technical PCR replicates (out of 16) in which the species was detected, summed across markers, plotted against year. Line indicates a per-site loess fit.

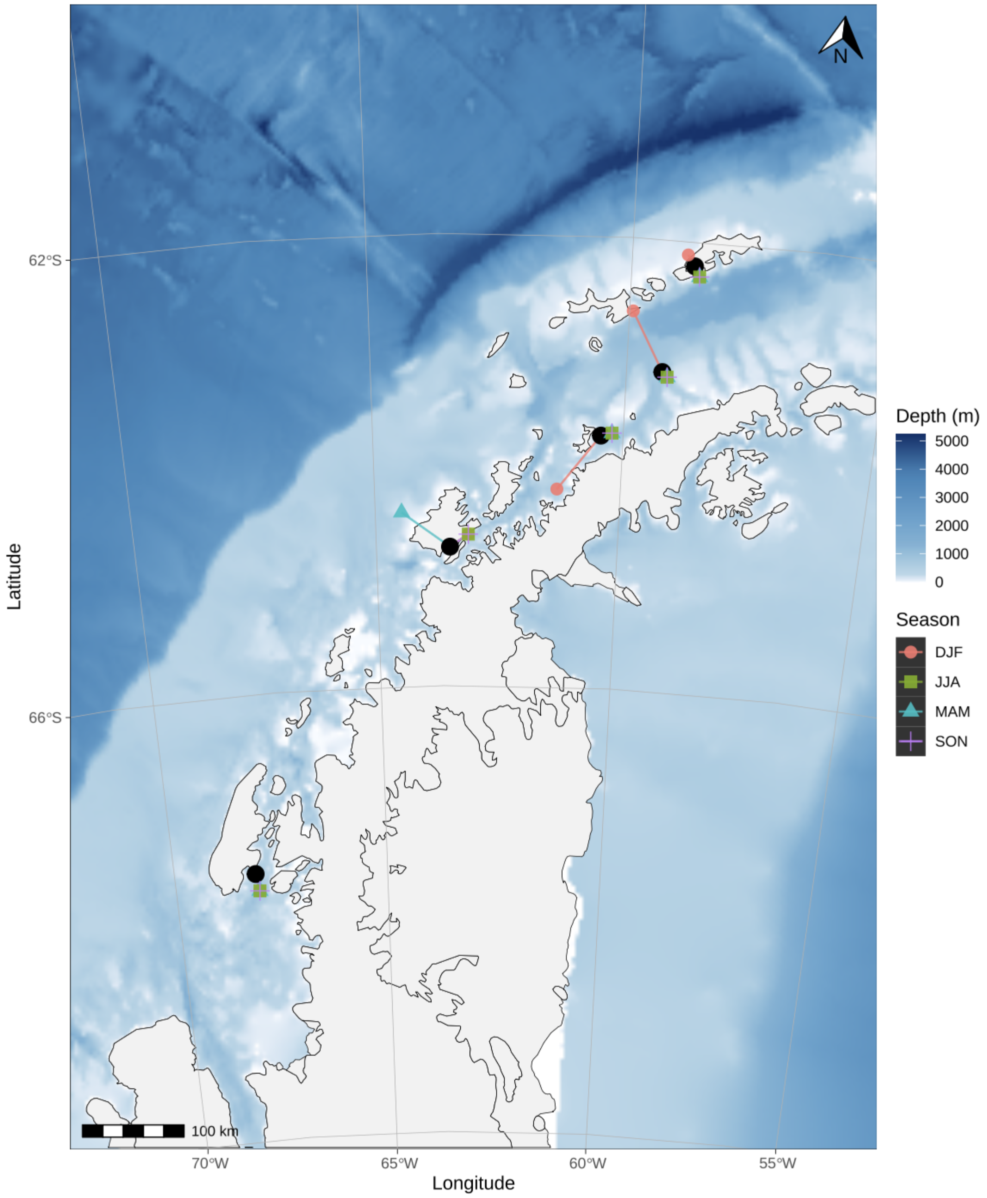


**Supplementary Figure 16:** Map of sampling locations (black dots) across the Western Antarctic Peninsula. For each sampling point the centre of the nearest available data point for sea ice concentration from Fogt et al. [4] is indicated by a coloured point for each of four seasons. The seasons are DJF (December, January, February), JJA (June, July, August), MAM (March, April, May) and SON (September, October, November).

**Supplementary Table 1: a** Mean count and standard deviation of read pairs per replicate from the five Western Antarctic Peninsula cores. The mean and s.d. are shown for raw demultiplexed reads, those surviving bioinformatic analyses (Post Bioinformatic), and those surviving final quality control and subject to ecological analyses (Final Dataset). The values are shown for each metabarcoding assay (18S rRNA V9, eukaryotes; 16S rRNA, mammals; 12S rRNA, vertebrates), and for experimental and no template (negative) controls. **b** Negative control sequencing results. Extraction and PCR blank replicates from both sequencing runs combined, per marker. Taxa are genus-level NCBI assignments.

## a

| | | Demultiplexed | Post Bioinformatics | Final Dataset |
|---|---|---|---|---|
| **18S** | **Experimental** | 247679.9 ± 214545.2 | 217275.1 ± 190959.5 | 216803.5 ± 190690.6 |
| | **Negative Controls** | 212.4 ± 1109.3 | 30.6 ± 55.8 | NA |
| **16S** | **Experimental** | 87198.7 ± 130662.9 | 74300.1 ± 107813 | 49114.5 ± 80723.3 |
| | **Negative Controls** | 32773.3 ± 64019.2 | 10538.8 ± 47101.9 | NA |
| **12S** | **Experimental** | 30690.6 ± 62622.7 | 32951.2 ± 60635.9 | 27837.1 ± 54742.1 |
| | **Negative Controls** | 5506.5 ± 20699.9 | 4399.5 ± 20311.4 | NA |

## b

| | Extraction blanks | PCR blanks | Median reads per replicate | Max reads per replicate | Total reads | Replicates with 1–99 reads | Replicates with >1,000 reads | Human reads (%) |
|---|---|---|---|---|---|---|---|---|
| **18S** | 56 | 73 | 9 | 402 | 3946 | 106 | 0 | 0.2 |
| **16S** | 52 | 73 | 4 | 342036 | 1317355 | 99 | 10 | 99.9 |
| **12S** | 44 | 54 | 2 | 163100 | 431152 | 69 | 9 | 59.5 |

| Top 5 most abundant non-human genera across all controls (total reads) | |
|---|---|
| **18S** | Chaetoceros (2203); Cryothecomonas (178); Ebria (58); Apatococcus (50); Botuliforma (38) |
| **16S** | Balaenoptera (340); Lobodon (66); Orcinus (25); Megaptera (21); Pan (19) |
| **12S** | Sus (99058); Canis (28719); Pleuragramma (125); Balaenoptera (119); Lobodon (37) |

**Supplementary Table 2:** Coefficients, standard errors and p-values from linear mixed models of within-sample diversity (Hill numbers of order 0, 1 and 2, computed on rarefied reads at ASV and genus level) against scaled Southern Annular Mode, ENSO, site sea ice concentration and summed whale biomass, with coring site as a random intercept (lme4). '*model*' column shows the response variable with taxonomic resolution (ASV or genus) and Hill number order. '*predictor'* gives the fixed effect: SAM_s (3-year mean Southern Annular Mode index), ENSO_s (3-year mean ENSO index), SeaIce_s (mean sea ice concentration at the coring site) and whale_s (summed whale biomass for the corresponding calendar year). '*estimate'* is the fixed-effect slope. '*SE'* is its standard error. '*df'* is the denominator degrees of freedom for the test of the estimate. '*t'* is the Wald t statistic. '*CI_lo'* and '*CI_hi'* are the lower and upper bounds of the 95% Wald confidence interval for the estimate. '*p_value'* is the two-sided p-value. '*sig'* is the corresponding significance code (** p < 0.01, * p < 0.05, . p < 0.1, blank p ≥ 0.1).

| model | predictor | estimate | SE | df | t | CI_lo | CI_hi | p_value | sig |
|---|---|---|---|---|---|---|---|---|---|
| ASV Q0 | SAM_s | 43.5433 | 15.9923 | 78.5404 | 2.7228 | 12.1989 | 74.8878 | 0.008 | ** |
| ASV Q0 | ENSO_s | -3.0948 | 13.1157 | 78.5144 | -0.236 | -28.8011 | 22.6116 | 0.8141 | |
| ASV Q0 | SeaIce_s | 2.3261 | 29.1756 | 79.6236 | 0.0797 | -54.8569 | 59.5092 | 0.9367 | |
| ASV Q0 | whale_s | 1.5371 | 15.203 | 77.9895 | 0.1011 | -28.2603 | 31.3344 | 0.9197 | |
| ASV Q1 | SAM_s | 0.9329 | 1.9863 | 78.6399 | 0.4697 | -2.9601 | 4.8259 | 0.6399 | |
| ASV Q1 | ENSO_s | -0.9111 | 1.6291 | 78.5987 | -0.5593 | -4.104 | 2.2818 | 0.5776 | |
| ASV Q1 | SeaIce_s | 0.002 | 3.5519 | 72.5282 | 0.0006 | -6.9596 | 6.9637 | 0.9995 | |
| ASV Q1 | whale_s | 2.9346 | 1.8899 | 77.8412 | 1.5528 | -0.7695 | 6.6387 | 0.1245 | |
| ASV Q2 | SAM_s | 0.3721 | 0.5568 | 79.0947 | 0.6682 | -0.7193 | 1.4634 | 0.5059 | |
| ASV Q2 | ENSO_s | -0.1221 | 0.4567 | 79.0269 | -0.2673 | -1.0172 | 0.773 | 0.7899 | |
| ASV Q2 | SeaIce_s | 0.3815 | 0.9596 | 56.0968 | 0.3976 | -1.4993 | 2.2623 | 0.6924 | |
| ASV Q2 | whale_s | 0.8155 | 0.5306 | 77.965 | 1.537 | -0.2244 | 1.8555 | 0.1283 | |
| Genus Q0 | SAM_s | 6.913 | 3.034 | 78.8808 | 2.2785 | 0.9665 | 12.8595 | 0.0254 | * |
| Genus Q0 | ENSO_s | -2.253 | 2.4883 | 78.8438 | -0.9054 | -7.1301 | 2.624 | 0.368 | |
| Genus Q0 | SeaIce_s | 3.7897 | 5.4351 | 73.9008 | 0.6973 | -6.8629 | 14.4423 | 0.4878 | |
| Genus Q0 | whale_s | 1.4645 | 2.8865 | 78.1584 | 0.5074 | -4.1929 | 7.122 | 0.6133 | |
| Genus Q1 | SAM_s | 0.4102 | 0.3032 | 78.8782 | 1.3527 | -0.1841 | 1.0045 | 0.18 | |
| Genus Q1 | ENSO_s | -0.1198 | 0.2487 | 78.8334 | -0.4817 | -0.6072 | 0.3676 | 0.6313 | |
| Genus Q1 | SeaIce_s | -0.0409 | 0.538 | 69.6399 | -0.0761 | -1.0955 | 1.0136 | 0.9396 | |
| Genus Q1 | whale_s | 0.5879 | 0.2886 | 78.0368 | 2.0369 | 0.0222 | 1.1535 | 0.045 | * |
| Genus Q2 | SAM_s | 0.251 | 0.1567 | 79.4094 | 1.6015 | -0.0562 | 0.5582 | 0.1132 | |
| Genus Q2 | ENSO_s | 0.0048 | 0.1286 | 79.3408 | 0.0371 | -0.2472 | 0.2567 | 0.9705 | |
| Genus Q2 | SeaIce_s | -0.0098 | 0.2678 | 54.0293 | -0.0367 | -0.5347 | 0.515 | 0.9709 | |
| Genus Q2 | whale_s | 0.2539 | 0.1494 | 78.303 | 1.6995 | -0.0389 | 0.5468 | 0.0932 | . |

**Supplementary Table 3:** Distance-based redundancy analysis (dbRDA) of community composition against scaled whale biomass, Southern Annular Mode, ENSO and site sea ice concentration, with coring site as a conditioning variable, fitted separately to the whole community and to metazoan and protist ASV subsets. Significance was assessed by permutation (100,000 permutations, restricted within sites). 'Group' gives the taxonomic subset analysed. 'Distance' gives the dissimilarity measure, either Bray–Curtis on proportional abundances or Jaccard on presence–absence data. 'Term' is either the overall constrained model or an individual predictor, tested marginally. 'Df' is the degrees of freedom, 'SumOfSqs' the constrained sum of squares and 'F_value' the pseudo-F statistic for that term. 'p_value' is the permutation p-value. 'R2_constrained' is the proportion of total inertia explained by the constrained axes, given once per model on the 'Overall model' row and left blank for individual terms.

| Group | Distance | Term | Df | SumOfSqs | F_value | p_value | R2_constrained |
|---|---|---|---|---|---|---|---|
| Global | Bray-Curtis | Overall model | 4 | 0.9349 | 2.855 | <0.001 | 0.128 |
| Global | Bray-Curtis | whale_s | 1 | 0.1344 | 1.642 | 0.029 | |
| Global | Bray-Curtis | SAM_s | 1 | 0.4409 | 5.385 | <0.001 | |
| Global | Bray-Curtis | ENSO_s | 1 | 0.0504 | 0.616 | 0.838 | |
| Global | Bray-Curtis | SeaIce_s | 1 | 0.0592 | 0.723 | 0.856 | |
| Global | Jaccard | Overall model | 4 | 1.9697 | 1.915 | <0.001 | 0.089 |
| Global | Jaccard | whale_s | 1 | 0.6509 | 2.531 | <0.001 | |
| Global | Jaccard | SAM_s | 1 | 0.5975 | 2.324 | <0.001 | |
| Global | Jaccard | ENSO_s | 1 | 0.2543 | 0.989 | 0.484 | |
| Global | Jaccard | SeaIce_s | 1 | 0.2543 | 0.989 | 0.364 | |
| Metazoa | Bray-Curtis | Overall model | 4 | 2.3023 | 3.504 | <0.001 | 0.152 |
| Metazoa | Bray-Curtis | whale_s | 1 | 0.4492 | 2.735 | 0.007 | |
| Metazoa | Bray-Curtis | SAM_s | 1 | 0.9392 | 5.717 | <0.001 | |
| Metazoa | Bray-Curtis | ENSO_s | 1 | 0.1809 | 1.101 | 0.294 | |
| Metazoa | Bray-Curtis | SeaIce_s | 1 | 0.2437 | 1.484 | 0.103 | |
| Metazoa | Jaccard | Overall model | 4 | 1.6163 | 1.502 | <0.001 | 0.072 |
| Metazoa | Jaccard | whale_s | 1 | 0.5445 | 2.024 | <0.001 | |
| Metazoa | Jaccard | SAM_s | 1 | 0.4807 | 1.787 | <0.001 | |
| Metazoa | Jaccard | ENSO_s | 1 | 0.2917 | 1.084 | 0.186 | |
| Metazoa | Jaccard | SeaIce_s | 1 | 0.2754 | 1.024 | 0.277 | |
| Protists | Bray-Curtis | Overall model | 4 | 0.5706 | 2.746 | <0.001 | 0.123 |
| Protists | Bray-Curtis | whale_s | 1 | 0.0811 | 1.560 | 0.056 | |
| Protists | Bray-Curtis | SAM_s | 1 | 0.2872 | 5.529 | 0.000 | |
| Protists | Bray-Curtis | ENSO_s | 1 | 0.0307 | 0.592 | 0.735 | |
| Protists | Bray-Curtis | SeaIce_s | 1 | 0.0203 | 0.391 | 0.968 | |
| Protists | Jaccard | Overall model | 4 | 1.8116 | 2.039 | <0.001 | 0.095 |
| Protists | Jaccard | whale_s | 1 | 0.6442 | 2.901 | <0.001 | |
| Protists | Jaccard | SAM_s | 1 | 0.5583 | 2.514 | <0.001 | |
| Protists | Jaccard | ENSO_s | 1 | 0.2179 | 0.981 | 0.510 | |
| Protists | Jaccard | SeaIce_s | 1 | 0.2132 | 0.960 | 0.519 | |

**Supplementary Table 4:** Mean posterior regression coefficients and posterior support for the direction of each effect (SAM, ENSO, sea ice concentration, whale biomass), and the proportion of total variance attributed to climate, whale biomass, the site random effect and the residual, for each of the 55 metazoan families in the abundance-based joint species distribution model (HMSC; centred-log-ratio-transformed relative abundances, normal distribution). 'taxon' gives the family, with † marking taxa resolved only to a higher rank. 'beta_SAM', 'beta_ENSO', 'beta_SeaIce' and 'beta_whale' are the posterior mean slopes for each scaled predictor. 'support_SAM', 'support_ENSO', 'support_SeaIce' and 'support_whale' are the posterior probabilities that the corresponding coefficient has the sign of its posterior mean. 'var_climate_pct', 'var_whaling_pct', 'var_site_pct' and 'var_residual_pct' are the percentages of total variance attributed to the climate predictors (SAM, ENSO and sea ice combined), whale biomass, the site random effect and the residual; variance proportions are the partitioned fraction of explained variance multiplied by each family's explanatory $R^2$, and the final row gives means across families. Explanatory $R^2$ for each taxon is 100 − var_residual_pct.

| Taxon | beta_SAM | support_SAM | beta_ENSO | support_ENSO | beta_SeaIce | support_SeaIce | beta_whale | support_whale | var_climate_pct | var_whaling_pct | var_site_pct | var_residual_pct |
|---|---|---|---|---|---|---|---|---|---|---|---|---|
| Salpidae | 0.053 | 0.711 | 0.006 | 0.531 | -0.057 | 0.756 | 0.802 | 1 | 2.2 | 50.6 | 0.6 | 46.6 |
| Ascidiidae | 0.118 | 0.864 | 0.058 | 0.744 | -0.27 | 0.994 | 0.737 | 1 | 7.8 | 34.8 | 7.6 | 49.8 |
| Edwardsiidae | 0.141 | 0.793 | -0.08 | 0.716 | 0.597 | 0.99 | 0.702 | 1 | 17.7 | 21.5 | 7.4 | 53.4 |
| Prayidae | 0.136 | 0.811 | -0.083 | 0.746 | 0.539 | 0.996 | 0.619 | 1 | 18.7 | 21.8 | 4.8 | 54.7 |
| Periphyllidae | -0.001 | 0.503 | 0 | 0.502 | -0.12 | 0.726 | 0.482 | 0.998 | 7.2 | 15.6 | 10 | 67.3 |
| Euphausiidae | 0.388 | 0.993 | 0.014 | 0.544 | 0.128 | 0.78 | 0.453 | 0.999 | 8.6 | 8.7 | 18.7 | 63.9 |
| Terebellidae | 0.213 | 0.917 | -0.035 | 0.613 | 0.305 | 0.968 | 0.372 | 0.994 | 9.1 | 7.8 | 19.7 | 63.5 |
| Maldanidae | -0.064 | 0.643 | 0.003 | 0.51 | 0.195 | 0.816 | 0.23 | 0.92 | 9.1 | 5.3 | 29.3 | 56.3 |
| Sternaspidae | 0.165 | 0.816 | -0.247 | 0.952 | 0.705 | 0.999 | 0.212 | 0.892 | 23 | 3 | 20.6 | 53.5 |
| Ampharetidae | -0.376 | 0.991 | 0.041 | 0.629 | -0.195 | 0.889 | 0.181 | 0.888 | 12.7 | 3.6 | 0.6 | 83.1 |
| Echiuridae | -0.14 | 0.803 | -0.082 | 0.732 | 0.644 | 0.998 | 0.157 | 0.849 | 26.3 | 2.7 | 23.1 | 47.9 |
| Oikopleuridae | -0.333 | 0.987 | -0.067 | 0.716 | -0.174 | 0.881 | 0.081 | 0.725 | 14.8 | 2.3 | 7.3 | 75.7 |
| Inversiulidae | -0.028 | 0.585 | 0.179 | 0.951 | -0.751 | 1 | 0.064 | 0.697 | 44.2 | 1.5 | 15.3 | 39 |
| Lysianassidae | 0.022 | 0.57 | 0.006 | 0.521 | -0.188 | 0.95 | 0.063 | 0.71 | 8.4 | 2 | 8.7 | 80.9 |
| Bugulidae | 0.006 | 0.515 | 0.003 | 0.51 | 0.296 | 0.972 | 0.053 | 0.644 | 6.9 | 1.2 | 1.7 | 90.1 |
| Cheilostomatida † | -0.046 | 0.609 | -0.117 | 0.801 | -0.384 | 0.983 | 0.024 | 0.557 | 15.1 | 1.8 | 9.3 | 73.7 |
| Tisbidae | 0.008 | 0.519 | 0.035 | 0.616 | -0.197 | 0.905 | 0.013 | 0.537 | 11.3 | 2.5 | 10.5 | 75.7 |
| Oithonidae | -0.111 | 0.803 | -0.072 | 0.757 | 0.216 | 0.937 | 0.001 | 0.508 | 10.2 | 1.6 | 37.6 | 50.6 |
| Ulmaridae | -0.186 | 0.914 | 0.042 | 0.646 | -0.179 | 0.89 | -0.011 | 0.536 | 14.7 | 2.6 | 14 | 68.6 |
| Cyclopoida † | 0.098 | 0.804 | 0.125 | 0.907 | 0.04 | 0.641 | -0.028 | 0.598 | 9 | 2 | 13.4 | 75.5 |
| Nacellidae | -0.131 | 0.844 | 0.015 | 0.556 | -0.729 | 1 | -0.04 | 0.629 | 37 | 1.1 | 30 | 31.8 |
| Orbiniidae | -0.006 | 0.519 | -0.036 | 0.626 | -0.153 | 0.864 | -0.04 | 0.624 | 9.1 | 2.4 | 7 | 81.6 |
| Yoldiidae | 0.063 | 0.674 | -0.062 | 0.711 | 0.033 | 0.595 | -0.063 | 0.688 | 4.6 | 1.7 | 4.4 | 89.3 |
| Rhopalonematidae | -0.042 | 0.633 | 0.174 | 0.956 | -0.29 | 0.984 | -0.071 | 0.729 | 18.9 | 2.8 | 16.7 | 61.5 |
| Pontogeneiidae | -0.168 | 0.907 | -0.097 | 0.824 | -0.298 | 0.993 | -0.076 | 0.734 | 10 | 1.2 | 0.5 | 88.2 |
| Miraciidae | 0.01 | 0.534 | 0.135 | 0.912 | -0.165 | 0.915 | -0.077 | 0.749 | 10.6 | 2.7 | 9.2 | 77.6 |
| Actinostolidae | -0.189 | 0.918 | -0.001 | 0.501 | 0.536 | 1 | -0.093 | 0.771 | 32.2 | 2 | 3.1 | 62.7 |
| Lineidae | -0.063 | 0.669 | -0.22 | 0.971 | 0.318 | 0.983 | -0.095 | 0.764 | 15.4 | 2.2 | 19.6 | 62.8 |
| Canthocamptidae | 0.003 | 0.511 | 0.149 | 0.919 | -0.004 | 0.509 | -0.103 | 0.799 | 6.4 | 2.3 | 7.2 | 84.1 |
| Monhysteridae | 0.428 | 0.994 | -0.054 | 0.659 | 0.317 | 0.971 | -0.12 | 0.784 | 18.6 | 2.7 | 12.6 | 66.1 |
| Callochitonidae | -0.029 | 0.593 | 0.12 | 0.885 | -0.065 | 0.713 | -0.125 | 0.859 | 5.6 | 2.7 | 6.9 | 84.7 |
| Lumbrineridae | -0.122 | 0.803 | 0.16 | 0.917 | -0.161 | 0.87 | -0.127 | 0.829 | 6.1 | 1.9 | 3.3 | 88.6 |
| Syllidae | -0.148 | 0.869 | -0.046 | 0.667 | -0.398 | 0.998 | -0.131 | 0.853 | 19.3 | 2.9 | 14.5 | 63.3 |
| Clavelinidae | 0.001 | 0.504 | -0.048 | 0.668 | -0.132 | 0.839 | -0.132 | 0.857 | 8.4 | 3.9 | 8.4 | 79.4 |
| Golfingiidae | -0.062 | 0.666 | -0.086 | 0.777 | 0.791 | 1 | -0.132 | 0.842 | 39.4 | 2 | 5.5 | 53.1 |
| Campanulariidae | -0.046 | 0.651 | -0.087 | 0.815 | 0.311 | 0.997 | -0.133 | 0.88 | 14.6 | 3.2 | 7.7 | 74.5 |
| Cirratulidae | -0.226 | 0.934 | 0.017 | 0.556 | -0.551 | 1 | -0.134 | 0.833 | 23.3 | 2.3 | 5 | 69.3 |
| Convolutidae | -0.04 | 0.622 | -0.015 | 0.557 | -0.07 | 0.709 | -0.137 | 0.87 | 4.5 | 2.7 | 2.2 | 90.7 |
| Leptolaimidae | 0.495 | 0.994 | 0.039 | 0.603 | 0.366 | 0.965 | -0.147 | 0.803 | 19.1 | 2.7 | 3.2 | 75 |
| Echinoderidae | 0.169 | 0.874 | -0.024 | 0.578 | -0.103 | 0.756 | -0.154 | 0.868 | 7.6 | 3.4 | 5.4 | 83.7 |
| Comesomatidae | 0.537 | 0.994 | 0.051 | 0.622 | 0.412 | 0.962 | -0.186 | 0.835 | 21.3 | 3.4 | 2.7 | 72.6 |
| Gadilidae | -0.043 | 0.651 | -0.051 | 0.714 | 0.157 | 0.924 | -0.206 | 0.974 | 8.1 | 6.6 | 14.1 | 71.2 |
| Oerstediidae | -0.056 | 0.708 | -0.053 | 0.738 | -0.271 | 0.992 | -0.209 | 0.984 | 19.3 | 9.5 | 20 | 51.2 |
| Camarodonta † | -0.142 | 0.857 | -0.055 | 0.694 | 0.583 | 1 | -0.22 | 0.961 | 24.8 | 3.7 | 3.8 | 67.8 |
| Cyamiidae | -0.042 | 0.633 | 0.031 | 0.622 | -0.392 | 0.998 | -0.224 | 0.971 | 24.6 | 8.5 | 9.4 | 57.5 |
| Opheliidae | -0.384 | 0.999 | -0.004 | 0.512 | -0.522 | 1 | -0.23 | 0.977 | 32 | 5 | 10 | 53.1 |
| Ectinosomatidae | -0.058 | 0.663 | -0.057 | 0.693 | 0.314 | 0.991 | -0.265 | 0.979 | 8 | 4.4 | 0.6 | 87.1 |
| Mopaliidae | -0.058 | 0.694 | 0.237 | 0.993 | -0.041 | 0.641 | -0.271 | 0.993 | 10.1 | 8.9 | 8.4 | 72.6 |
| Lacernidae | -0.071 | 0.714 | 0.182 | 0.96 | -0.449 | 1 | -0.272 | 0.988 | 18.8 | 6.6 | 7.3 | 67.2 |
| Iphimediidae | -0.25 | 0.985 | 0.144 | 0.94 | -0.397 | 1 | -0.287 | 0.996 | 19.5 | 7.8 | 6.9 | 65.9 |
| Spionidae | -0.092 | 0.797 | 0.033 | 0.642 | -0.06 | 0.711 | -0.289 | 0.997 | 4.9 | 9.9 | 14.1 | 71.1 |
| Thyasiridae | -0.156 | 0.899 | -0.141 | 0.924 | -0.155 | 0.911 | -0.294 | 0.993 | 8.7 | 7.5 | 2 | 81.8 |
| Nephtheidae | 0.136 | 0.838 | 0.099 | 0.812 | 0.322 | 0.989 | -0.302 | 0.99 | 18 | 10.8 | 5.6 | 65.6 |
| Pleustidae | -0.258 | 0.989 | 0.028 | 0.621 | -0.436 | 1 | -0.323 | 0.998 | 22.5 | 9.9 | 3 | 64.6 |
| Entoprocta † | -0.463 | 1 | 0.025 | 0.61 | -0.228 | 0.979 | -0.384 | 1 | 21.2 | 12.2 | 4.9 | 61.7 |
| Mean | -0.026 | 0.769 | 0.004 | 0.716 | -0.008 | 0.906 | -0.016 | 0.845 | 15.45 | 6.48 | 9.916 | 68.16 |

**Supplementary Table 5:** Mean posterior regression coefficients and posterior support for the direction of each effect (SAM, ENSO, sea ice concentration, whale biomass), and the proportion of total variance attributed to climate, whale biomass, the site random effect and the residual, for each of the 53 subdivisions in the abundance-based joint species distribution model (HMSC; centred-log-ratio-transformed relative abundances, normal distribution). 'taxon' gives the PR2 subdivision, including prokaryotic groups recovered as off-target amplification. 'beta_SAM', 'beta_ENSO', 'beta_SeaIce' and 'beta_whale' are the posterior mean slopes for each scaled predictor. 'support_SAM', 'support_ENSO', 'support_SeaIce' and 'support_whale' are the posterior probabilities that the corresponding coefficient has the sign of its posterior mean, so values approaching 1 indicate strong support and values near 0.5 indicate none. 'var_climate_pct', 'var_whaling_pct', 'var_site_pct' and 'var_residual_pct' are the percentages of total variance attributed to the climate predictors (SAM, ENSO and sea ice combined), whale biomass, the site random effect and the residual; variance proportions are the partitioned fraction of explained variance multiplied by each taxon's explanatory $R^2$, and the final row gives means across taxa. Explanatory $R^2$ for each taxon is 100 − var_residual_pct.

| taxon | beta_SAM | support_SAM | beta_ENSO | support_ENSO | beta_SeaIce | support_SeaIce | beta_whale | support_whale | var_climate_pct | var_whaling_pct | var_site_pct | var_residual_pct |
|---|---|---|---|---|---|---|---|---|---|---|---|---|
| Cryptophyta | 0.293 | 0.973 | -0.028 | 0.598 | 0.107 | 0.757 | 0.638 | 1 | 7.9 | 24 | 26.5 | 41.6 |
| Radiolaria | 0.216 | 0.964 | -0.028 | 0.618 | 0.154 | 0.885 | 0.635 | 1 | 7.7 | 33.4 | 24.1 | 34.8 |
| Picozoa_subdivision_1 | 0.24 | 0.939 | -0.01 | 0.533 | 0.034 | 0.601 | 0.378 | 0.994 | 7.2 | 10.1 | 15.6 | 67.1 |
| Haptophyta | 0.038 | 0.624 | -0.086 | 0.827 | 0.115 | 0.859 | 0.365 | 0.999 | 7 | 18.1 | 17.8 | 57.2 |
| Eurhodophytina | 0.208 | 0.882 | 0.032 | 0.598 | -0.58 | 0.999 | 0.326 | 0.97 | 26.9 | 7.3 | 28.7 | 37.1 |
| Euglenozoa | 0.322 | 0.986 | 0.026 | 0.6 | 0.271 | 0.984 | 0.306 | 0.984 | 11.4 | 5.7 | 1.5 | 81.4 |
| Chlorophyta | 0.179 | 0.999 | -0.074 | 0.943 | 0.117 | 0.976 | 0.305 | 1 | 14.8 | 29.1 | 12.8 | 43.3 |
| Metazoa | 0.366 | 1 | -0.088 | 0.911 | 0.073 | 0.839 | 0.302 | 1 | 18.7 | 12.1 | 3.1 | 66.1 |
| Choanoflagellata | 0.145 | 0.806 | -0.109 | 0.816 | 0.13 | 0.815 | 0.234 | 0.927 | 7.7 | 5.2 | 17.1 | 69.9 |
| Tubulinea | 0.055 | 0.625 | -0.03 | 0.593 | 0.089 | 0.714 | 0.229 | 0.911 | 6.7 | 5.5 | 7.1 | 80.7 |
| Planctomycetes | 0.228 | 0.986 | -0.036 | 0.675 | -0.13 | 0.924 | 0.216 | 0.984 | 9.4 | 4.9 | 1.1 | 84.7 |
| Breviatea | -0.127 | 0.893 | 0.104 | 0.9 | -0.104 | 0.854 | 0.215 | 0.983 | 12.8 | 12.2 | 8.1 | 66.9 |
| Acidobacteria | -0.371 | 0.978 | 0.065 | 0.691 | -0.1 | 0.744 | 0.193 | 0.869 | 11 | 4 | 1.9 | 83.1 |
| Apicomplexa | 0.373 | 1 | 0.043 | 0.697 | -0.154 | 0.946 | 0.164 | 0.944 | 16.9 | 2.7 | 0.9 | 79.5 |
| Dinoflagellata | -0.051 | 0.801 | 0.039 | 0.782 | -0.041 | 0.71 | 0.159 | 0.996 | 11.6 | 19 | 36.5 | 32.9 |
| Bigyra | -0.014 | 0.578 | -0.009 | 0.561 | 0.146 | 0.913 | 0.144 | 0.981 | 7.2 | 5.1 | 61.5 | 26.1 |
| Foraminifera | 0.483 | 0.995 | -0.026 | 0.576 | 0.343 | 0.979 | 0.142 | 0.793 | 18.3 | 2.3 | 2.6 | 76.8 |
| Discosea | 0.689 | 1 | -0.213 | 0.959 | 0.67 | 1 | 0.116 | 0.767 | 40.3 | 1.7 | 2.2 | 55.8 |
| Proteobacteria | -0.086 | 0.795 | 0.142 | 0.954 | -0.092 | 0.836 | 0.113 | 0.87 | 8.3 | 3.4 | 1.4 | 86.9 |
| Alveolata | -0.356 | 0.993 | 0.022 | 0.581 | 0.014 | 0.545 | 0.112 | 0.792 | 12.6 | 2.8 | 1.5 | 83.1 |
| Fungi | 0.19 | 0.88 | 0.054 | 0.674 | -0.025 | 0.574 | 0.067 | 0.671 | 8.5 | 2.3 | 7.5 | 81.6 |
| Apusomonada | 0.144 | 0.869 | 0.11 | 0.869 | 0.226 | 0.976 | 0.058 | 0.676 | 15.5 | 2.3 | 9.3 | 72.8 |
| Epsilonbacteraeota | -0.451 | 0.967 | 0.021 | 0.553 | -0.069 | 0.628 | 0.034 | 0.56 | 12.8 | 2.5 | 15.4 | 69.3 |
| Metamonada | -0.18 | 0.921 | 0.018 | 0.576 | -0.039 | 0.636 | -0.008 | 0.526 | 11.4 | 2.8 | 14.4 | 71.5 |
| Euryarchaeota | -0.93 | 1 | 0.108 | 0.764 | -0.216 | 0.876 | -0.013 | 0.526 | 36.3 | 1.8 | 11.9 | 50 |
| Fornicata | -0.453 | 0.991 | 0.006 | 0.516 | -0.277 | 0.938 | -0.029 | 0.567 | 14.6 | 1.7 | 50.1 | 33.7 |
| Picozoa_subdivision_2 | 0.112 | 0.761 | -0.066 | 0.716 | -0.045 | 0.629 | -0.074 | 0.688 | 8.8 | 3.3 | 10.8 | 77.1 |
| Thaumarchaeota | 0.434 | 0.991 | -0.054 | 0.663 | 0.412 | 0.995 | -0.083 | 0.691 | 23.2 | 2.6 | 7.3 | 66.9 |
| Telonemia | 0.079 | 0.699 | -0.146 | 0.902 | 0.305 | 0.986 | -0.087 | 0.723 | 16 | 3.1 | 18.4 | 62.5 |
| Opisthokonta | -0.217 | 0.966 | -0.001 | 0.504 | 0.073 | 0.759 | -0.092 | 0.791 | 14.1 | 3.2 | 26.6 | 56.1 |
| Firmicutes | -0.163 | 0.846 | -0.043 | 0.644 | -0.294 | 0.974 | -0.093 | 0.728 | 12.1 | 2.3 | 49.7 | 35.9 |
| Discoba | -0.192 | 0.884 | -0.035 | 0.619 | 0.244 | 0.949 | -0.104 | 0.755 | 13.6 | 2.7 | 12.3 | 71.5 |
| Verrucomicrobia | 0.515 | 0.997 | -0.021 | 0.569 | -0.182 | 0.882 | -0.128 | 0.776 | 24.2 | 3.4 | 1.8 | 70.5 |
| Nitrospirae | 0.33 | 0.964 | -0.149 | 0.875 | -0.123 | 0.78 | -0.134 | 0.784 | 16.1 | 4 | 10.2 | 69.7 |
| Perkinsea | -0.05 | 0.65 | -0.075 | 0.774 | 0.245 | 0.98 | -0.136 | 0.86 | 14.4 | 4.3 | 19.4 | 62 |
| Nitrospinae | 0.11 | 0.843 | -0.045 | 0.699 | 0.24 | 0.99 | -0.143 | 0.911 | 16.9 | 6 | 28.3 | 48.8 |
| Prasinodermophyta | -0.264 | 0.96 | 0.081 | 0.758 | -0.493 | 1 | -0.153 | 0.858 | 26.6 | 3.5 | 13.8 | 56.1 |
| Kiritimatiellaeota | -0.177 | 0.89 | -0.054 | 0.694 | 0.053 | 0.665 | -0.169 | 0.89 | 4.9 | 2.3 | 1.8 | 90.9 |
| Ancyromonadida | -0.091 | 0.783 | 0.051 | 0.716 | -0.054 | 0.699 | -0.175 | 0.941 | 7.3 | 6.2 | 24.8 | 61.8 |
| Centroplasthelida | -0.069 | 0.678 | -0.046 | 0.66 | 0.077 | 0.729 | -0.179 | 0.896 | 4.1 | 2.8 | 2.3 | 90.9 |
| Ichthyosporea | 0.163 | 0.974 | -0.048 | 0.763 | 0.095 | 0.887 | -0.2 | 0.993 | 15.8 | 14.8 | 11.7 | 57.8 |
| Cryptophyta:nucl | -0.18 | 0.891 | 0.14 | 0.897 | -0.175 | 0.903 | -0.218 | 0.939 | 11.4 | 6.1 | 3.6 | 78.9 |
| Chrompodellids | -0.141 | 0.883 | 0.063 | 0.752 | 0.167 | 0.845 | -0.235 | 0.981 | 3.4 | 2.2 | 69.7 | 24.7 |
| Gyrista | -0.334 | 0.999 | 0.041 | 0.704 | -0.226 | 0.986 | -0.242 | 0.994 | 15 | 5.8 | 59.1 | 20.1 |
| Rhodophyta | -0.212 | 0.937 | -0.055 | 0.703 | -0.291 | 0.986 | -0.242 | 0.967 | 16.2 | 6.7 | 6.9 | 70.1 |
| Hydrogenedentes | -0.272 | 0.999 | -0.01 | 0.561 | -0.161 | 0.975 | -0.251 | 0.999 | 16.9 | 10.6 | 41.5 | 31.1 |
| Nebulidia | -0.186 | 0.973 | 0.021 | 0.606 | -0.236 | 0.99 | -0.261 | 0.997 | 16.6 | 12.2 | 27.6 | 43.6 |
| Ciliophora | 0.03 | 0.581 | 0.052 | 0.681 | -0.072 | 0.712 | -0.27 | 0.97 | 4.2 | 5.4 | 0.6 | 89.8 |
| Streptophyta | -0.229 | 0.89 | 0.003 | 0.511 | 0.114 | 0.762 | -0.306 | 0.958 | 7.9 | 5.9 | 16.6 | 69.6 |
| Hemimastigophora | -0.334 | 0.993 | 0.019 | 0.573 | -0.196 | 0.951 | -0.345 | 0.996 | 13.8 | 10.4 | 15.7 | 60.1 |
| Evosea | 0.289 | 0.93 | 0.17 | 0.887 | 0.225 | 0.91 | -0.346 | 0.968 | 13.7 | 8.3 | 3.4 | 74.6 |
| Dadabacteria | 0.122 | 0.762 | 0.087 | 0.75 | -0.088 | 0.705 | -0.355 | 0.984 | 10.3 | 13.3 | 11.8 | 64.7 |
| Cercozoa | -0.001 | 0.5 | -0.035 | 0.746 | -0.121 | 0.968 | -0.485 | 1 | 5.8 | 49.8 | 10.8 | 33.6 |
| Mean | 0.004 | 0.881 | -0.002 | 0.704 | 0.003 | 0.851 | -0.002 | 0.874 | 13.525 | 7.985 | 16.738 | 61.76 |

**Supplementary Table 6:** Variation partitioning of community composition among three explanatory sets — climate (scaled Southern Annular Mode, ENSO and site sea ice concentration), scaled whale biomass, and coring site — fitted separately to the whole community and to metazoan and protist ASV subsets (vegan). 'Group' gives the taxonomic subset analysed. 'Distance' gives the dissimilarity measure, either Bray–Curtis on proportional abundances or Jaccard on presence–absence data. 'Fraction' identifies the component of variation: the first three rows of each model are the unique contributions of each explanatory set conditional on the other two, the next four are the variation shared between sets and cannot be uniquely attributed, and the final row is the unexplained residual. 'Df' is the number of explanatory variables in each unique fraction, and is undefined for shared and residual fractions. 'Adj_R2' is the adjusted coefficient of determination for each fraction; adjusted values are unbiased and additive, so fractions sum to one within a model, but can take small negative values where a set explains less variation than expected by chance, which should be interpreted as zero.

| Group | Distance | Fraction | Df | Adj_R2 |
|---|---|---|---|---|
| Global | Bray-Curtis | Climate (SAM+ENSO+SeaIce) \| Whale + Site | 3 | 0.025 |
| Global | Bray-Curtis | Whale \| Climate + Site | 1 | 0.004 |
| Global | Bray-Curtis | Site \| Climate + Whale | 4 | 0.378 |
| Global | Bray-Curtis | Shared: Climate & Whale | | 0.016 |
| Global | Bray-Curtis | Shared: Whale & Site | | -0.006 |
| Global | Bray-Curtis | Shared: Climate & Site | | 0.066 |
| Global | Bray-Curtis | Shared: Climate & Whale & Site | | 0.020 |
| Global | Bray-Curtis | Residual | | 0.497 |
| Global | Jaccard | Climate (SAM+ENSO+SeaIce) \| Whale + Site | 3 | 0.013 |
| Global | Jaccard | Whale \| Climate + Site | 1 | 0.015 |
| Global | Jaccard | Site \| Climate + Whale | 4 | 0.171 |
| Global | Jaccard | Shared: Climate & Whale | | 0.006 |
| Global | Jaccard | Shared: Whale & Site | | -0.003 |
| Global | Jaccard | Shared: Climate & Site | | 0.031 |
| Global | Jaccard | Shared: Climate & Whale & Site | | 0.007 |
| Global | Jaccard | Residual | | 0.761 |
| Metazoa | Bray-Curtis | Climate (SAM+ENSO+SeaIce) \| Whale + Site | 3 | 0.037 |
| Metazoa | Bray-Curtis | Whale \| Climate + Site | 1 | 0.014 |
| Metazoa | Bray-Curtis | Site \| Climate + Whale | 4 | 0.221 |
| Metazoa | Bray-Curtis | Shared: Climate & Whale | | 0.025 |
| Metazoa | Bray-Curtis | Shared: Whale & Site | | -0.003 |
| Metazoa | Bray-Curtis | Shared: Climate & Site | | 0.076 |
| Metazoa | Bray-Curtis | Shared: Climate & Whale & Site | | 0.009 |
| Metazoa | Bray-Curtis | Residual | | 0.622 |
| Metazoa | Jaccard | Climate (SAM+ENSO+SeaIce) \| Whale + Site | 3 | 0.009 |
| Metazoa | Jaccard | Whale \| Climate + Site | 1 | 0.010 |
| Metazoa | Jaccard | Site \| Climate + Whale | 4 | 0.176 |
| Metazoa | Jaccard | Shared: Climate & Whale | | 0.000 |
| Metazoa | Jaccard | Shared: Whale & Site | | -0.002 |
| Metazoa | Jaccard | Shared: Climate & Site | | 0.026 |
| Metazoa | Jaccard | Shared: Climate & Whale & Site | | 0.007 |
| Metazoa | Jaccard | Residual | | 0.775 |
| Protists | Bray-Curtis | Climate (SAM+ENSO+SeaIce) \| Whale + Site | 3 | 0.021 |
| Protists | Bray-Curtis | Whale \| Climate + Site | 1 | 0.003 |
| Protists | Bray-Curtis | Site \| Climate + Whale | 4 | 0.431 |
| Protists | Bray-Curtis | Shared: Climate & Whale | | 0.013 |
| Protists | Bray-Curtis | Shared: Whale & Site | | -0.006 |
| Protists | Bray-Curtis | Shared: Climate & Site | | 0.069 |
| Protists | Bray-Curtis | Shared: Climate & Whale & Site | | 0.031 |
| Protists | Bray-Curtis | Residual | | 0.438 |
| Protists | Jaccard | Climate (SAM+ENSO+SeaIce) \| Whale + Site | 3 | 0.014 |
| Protists | Jaccard | Whale \| Climate + Site | 1 | 0.017 |
| Protists | Jaccard | Site \| Climate + Whale | 4 | 0.206 |
| Protists | Jaccard | Shared: Climate & Whale | | 0.005 |
| Protists | Jaccard | Shared: Whale & Site | | -0.004 |
| Protists | Jaccard | Shared: Climate & Site | | 0.031 |
| Protists | Jaccard | Shared: Climate & Whale & Site | | 0.009 |
| Protists | Jaccard | Residual | | 0.721 |

**Supplementary Table 7:** Contrasts in HMSC regression coefficients between water-column-associated taxa (pelagic water-column feeders and benthic suspension feeders) and all other metazoan families, for each predictor and for both the abundance-based (CLR) and occurrence-based (presence–absence) models. All modelled families are included, with no filtering on posterior support. 'model' identifies the HMSC model, 'predictor' the fixed effect, and 'n_water' and 'n_other' the number of families in each guild grouping. 'diff_mean' is the posterior mean difference in coefficient between the two groupings (water-column minus other), computed within each posterior sample, with 'diff_lo95' and 'diff_hi95' giving the bounds of the 95% credible interval. 'P_diff_gt0' is the posterior probability that this difference is positive. 'superiority' is the probability that a randomly chosen water-column family has a larger coefficient than a randomly chosen other family, again computed within posterior samples, with 'sup_lo95' and 'sup_hi95' its 95% credible interval; 0.5 indicates no difference. 'wilcoxon_W' and 'wilcoxon_p' are the statistic and p-value from a Wilcoxon rank-sum test on the posterior mean coefficients, given as a frequentist companion to the Bayesian contrasts.

| model | predictor | n_water | n_other | diff_mean | diff_lo95 | diff_hi95 | P_diff_gt0 | superiority | sup_lo95 | sup_hi95 | wilcoxon_W | wilcoxon_p |
|---|---|---|---|---|---|---|---|---|---|---|---|---|
| Metazoa CLR | SAM | 20 | 35 | -0.004 | -0.083 | 0.074 | 0.459 | 0.519 | 0.411 | 0.624 | 382 | 0.585 |
| Metazoa CLR | ENSO | 20 | 35 | 0.014 | -0.049 | 0.078 | 0.659 | 0.524 | 0.4 | 0.651 | 366 | 0.788 |
| Metazoa CLR | SeaIce | 20 | 35 | -0.025 | -0.106 | 0.056 | 0.275 | 0.483 | 0.42 | 0.547 | 331 | 0.748 |
| Metazoa CLR | whale | 20 | 35 | 0.214 | 0.14 | 0.286 | 1 | 0.655 | 0.57 | 0.736 | 473 | 0.031 |
| Metazoa PA | SAM | 19 | 35 | -0.059 | -0.166 | 0.043 | 0.13 | 0.444 | 0.311 | 0.576 | 289 | 0.44 |
| Metazoa PA | ENSO | 19 | 35 | 0.04 | -0.045 | 0.126 | 0.822 | 0.558 | 0.424 | 0.69 | 394 | 0.272 |
| Metazoa PA | SeaIce | 19 | 35 | -0.067 | -0.166 | 0.032 | 0.093 | 0.434 | 0.335 | 0.537 | 281 | 0.359 |
| Metazoa PA | whale | 19 | 35 | 0.169 | 0.054 | 0.289 | 0.998 | 0.644 | 0.555 | 0.729 | 444 | 0.044 |

**Supplementary Table 8:** Mean posterior regression coefficients and posterior support for the direction of each effect (SAM, ENSO, sea ice concentration, whale biomass), and the proportion of total variance attributed to climate, whale biomass, the site random effect and the residual, for each of the 54 metazoan families in the occurrence-based joint species distribution model (HMSC; presence–absence data, probit distribution). 'taxon' gives the family, with † marking taxa resolved only to a higher rank. 'beta_SAM', 'beta_ENSO', 'beta_SeaIce' and 'beta_whale' are the posterior mean slopes for each scaled predictor, on the probit scale. 'support_SAM', 'support_ENSO', 'support_SeaIce' and 'support_whale' are the posterior probabilities that the corresponding coefficient has the sign of its posterior mean, so values approaching 1 indicate strong support and values near 0.5 indicate none. 'var_climate_pct', 'var_whaling_pct', 'var_site_pct' and 'var_residual_pct' are the percentages of total variance attributed to the climate predictors (SAM, ENSO and sea ice combined), whale biomass, the site random effect and the residual; variance proportions are the partitioned fraction of explained variance multiplied by each family's explanatory $R^2$, and the final row gives means across families. Explanatory power for this probit model is Tjur $R^2$, equal to 100 − var_residual_pct for each family.

| taxon | beta_SAM | support_SAM | beta_ENSO | support_ENSO | beta_SeaIce | support_SeaIce | beta_whale | support_whale | var_climate_pct | var_whaling_pct | var_site_pct | var_residual_pct |
|---|---|---|---|---|---|---|---|---|---|---|---|---|
| Ascidiidae | 0.081 | 0.625 | 0.102 | 0.691 | 0.108 | 0.64 | 0.671 | 0.988 | 0.9 | 1.8 | 0.1 | 97.2 |
| Prayidae | 0.125 | 0.755 | -0.039 | 0.606 | 0.256 | 0.885 | 0.537 | 0.995 | 2.7 | 4.7 | 0.5 | 92.1 |
| Euphausiidae | 0.223 | 0.897 | -0.027 | 0.576 | 0.047 | 0.577 | 0.457 | 0.997 | 6.1 | 9.7 | 7.3 | 76.9 |
| Edwardsiidae | 0.117 | 0.762 | -0.124 | 0.815 | 0.205 | 0.843 | 0.419 | 0.995 | 7.9 | 10.7 | 7.7 | 73.7 |
| Terebellidae | 0.279 | 0.954 | -0.016 | 0.548 | 0.233 | 0.874 | 0.395 | 0.993 | 11.7 | 10.5 | 12.4 | 65.4 |
| Periphyllidae | 0.059 | 0.649 | -0.039 | 0.625 | 0.043 | 0.619 | 0.382 | 0.991 | 2.3 | 4.8 | 0.8 | 92.2 |
| Inversiulidae | 0.167 | 0.811 | 0.192 | 0.886 | -0.423 | 0.991 | 0.318 | 0.961 | 28.1 | 10.5 | 24.6 | 36.8 |
| Maldanidae | 0.034 | 0.587 | 0.02 | 0.557 | 0.219 | 0.91 | 0.313 | 0.982 | 11.4 | 11.2 | 14.3 | 63.1 |
| Ampharetidae | -0.215 | 0.928 | 0.001 | 0.501 | 0.035 | 0.599 | 0.259 | 0.961 | 9.2 | 7.5 | 3.4 | 80 |
| Sternaspidae | -0.029 | 0.572 | -0.275 | 0.984 | 0.674 | 1 | 0.241 | 0.946 | 26 | 4 | 0.9 | 69.1 |
| Oithonidae | 0.044 | 0.601 | -0.053 | 0.648 | 0.295 | 0.946 | 0.219 | 0.914 | 18.4 | 8.5 | 25.9 | 47.2 |
| Oikopleuridae | -0.164 | 0.842 | 0.001 | 0.5 | -0.021 | 0.557 | 0.176 | 0.877 | 12.6 | 7.3 | 17.3 | 62.8 |
| Cheilostomatida † | 0.1 | 0.743 | -0.228 | 0.964 | -0.03 | 0.571 | 0.157 | 0.858 | 5 | 1.8 | 1.1 | 92.1 |
| Ulmaridae | -0.152 | 0.826 | -0.064 | 0.692 | 0.199 | 0.91 | 0.149 | 0.84 | 13.7 | 4.8 | 8.6 | 72.9 |
| Echiuridae | -0.006 | 0.516 | -0.042 | 0.628 | 0.228 | 0.915 | 0.145 | 0.83 | 13.5 | 5.2 | 14.7 | 66.7 |
| Rhopalonematidae | 0.092 | 0.709 | 0.072 | 0.69 | 0.176 | 0.877 | 0.125 | 0.776 | 14 | 4.9 | 21.7 | 59.3 |
| Nacellidae | -0.004 | 0.507 | 0.045 | 0.616 | -0.345 | 0.984 | 0.114 | 0.743 | 23.5 | 5.3 | 31.5 | 39.7 |
| Bugulidae | 0.051 | 0.63 | -0.009 | 0.527 | 0.335 | 0.982 | 0.105 | 0.763 | 16.3 | 3.5 | 9.6 | 70.6 |
| Tisbidae | 0.183 | 0.874 | 0.035 | 0.598 | 0.022 | 0.56 | 0.064 | 0.656 | 14.3 | 4.2 | 23.9 | 57.5 |
| Pontogeneiidae | -0.048 | 0.614 | -0.154 | 0.875 | -0.041 | 0.612 | 0.054 | 0.632 | 11.6 | 3.6 | 13 | 71.8 |
| Lineidae | 0.094 | 0.715 | -0.326 | 0.992 | 0.43 | 0.994 | 0.052 | 0.62 | 28.8 | 3.1 | 16.7 | 51.4 |
| Orbiniidae | 0.148 | 0.812 | -0.049 | 0.636 | 0.072 | 0.693 | 0.049 | 0.615 | 12.5 | 4.2 | 23.6 | 59.8 |
| Yoldiidae | 0.203 | 0.889 | -0.198 | 0.92 | 0.298 | 0.985 | 0.038 | 0.59 | 12.6 | 2 | 3.7 | 81.7 |
| Cirratulidae | -0.111 | 0.734 | -0.052 | 0.644 | -0.172 | 0.822 | 0.028 | 0.57 | 6.5 | 1.6 | 2 | 89.8 |
| Cyclopoida † | 0.267 | 0.935 | 0.018 | 0.544 | 0.266 | 0.969 | 0.022 | 0.552 | 12.4 | 2.2 | 4 | 81.4 |
| Canthocamptidae | 0.162 | 0.826 | 0.049 | 0.627 | 0.122 | 0.805 | 0.004 | 0.516 | 7.4 | 2.2 | 4.3 | 86.1 |
| Lumbrineridae | 0.032 | 0.577 | 0.055 | 0.642 | 0.168 | 0.885 | 0.003 | 0.51 | 7.8 | 2.3 | 6.4 | 83.5 |
| Syllidae | 0.07 | 0.656 | -0.164 | 0.866 | -0.035 | 0.589 | -0.002 | 0.502 | 15 | 4.7 | 35.2 | 45.1 |
| Lysianassidae | 0.103 | 0.709 | -0.107 | 0.749 | 0.172 | 0.865 | -0.023 | 0.546 | 6.6 | 2.1 | 7.5 | 83.9 |
| Clavelinidae | 0.144 | 0.802 | -0.158 | 0.865 | 0.019 | 0.556 | -0.06 | 0.625 | 6.6 | 2.2 | 3.4 | 87.8 |
| Golfingiidae | 0.153 | 0.818 | -0.143 | 0.847 | 0.508 | 0.998 | -0.073 | 0.666 | 28.6 | 3.1 | 13.3 | 55 |
| Callochitonidae | 0.108 | 0.735 | 0.024 | 0.561 | 0.204 | 0.921 | -0.079 | 0.662 | 8.4 | 2.6 | 5.7 | 83.3 |
| Convolutidae | -0.028 | 0.563 | -0.129 | 0.826 | 0.208 | 0.927 | -0.08 | 0.667 | 13.8 | 3.9 | 18.2 | 64.1 |
| Miraciidae | 0.168 | 0.825 | 0.079 | 0.692 | 0.118 | 0.783 | -0.107 | 0.704 | 8.8 | 3.3 | 8.6 | 79.4 |
| Monhysteridae | 0.393 | 0.985 | -0.2 | 0.891 | 0.449 | 0.998 | -0.129 | 0.735 | 14.7 | 2.2 | 0.4 | 82.7 |
| Echinoderidae | 0.22 | 0.893 | -0.199 | 0.904 | 0.124 | 0.784 | -0.141 | 0.759 | 7.8 | 2.7 | 3.5 | 85.9 |
| Opheliidae | -0.129 | 0.763 | -0.118 | 0.785 | -0.028 | 0.572 | -0.141 | 0.764 | 12.2 | 5.2 | 31.1 | 51.5 |
| Actinostolidae | -0.122 | 0.731 | -0.133 | 0.793 | 0.62 | 0.999 | -0.154 | 0.764 | 30.1 | 3.7 | 16.3 | 49.9 |
| Iphimediidae | -0.191 | 0.839 | 0.061 | 0.656 | -0.271 | 0.921 | -0.173 | 0.809 | 11.2 | 3.9 | 5.8 | 79.2 |
| Ectinosomatidae | 0.026 | 0.563 | -0.148 | 0.857 | 0.312 | 0.987 | -0.198 | 0.863 | 15.5 | 5.6 | 10.5 | 68.4 |
| Campanulariidae | 0.086 | 0.672 | -0.303 | 0.966 | 0.513 | 0.999 | -0.211 | 0.835 | 20 | 4.1 | 5.5 | 70.4 |
| Nephtheidae | 0.211 | 0.908 | -0.017 | 0.556 | 0.285 | 0.977 | -0.236 | 0.922 | 14.2 | 6.6 | 3.4 | 75.9 |
| Camarodonta † | 0.135 | 0.757 | -0.126 | 0.792 | 0.581 | 1 | -0.272 | 0.908 | 26.5 | 6.8 | 20.9 | 45.7 |
| Leptolaimidae | 0.581 | 0.999 | -0.099 | 0.729 | 0.327 | 0.982 | -0.273 | 0.873 | 17.9 | 4.7 | 0.5 | 76.8 |
| Comesomatidae | 0.595 | 1 | -0.067 | 0.673 | 0.428 | 0.999 | -0.303 | 0.917 | 23 | 5.8 | 0.6 | 70.6 |
| Gadilidae | 0.085 | 0.667 | -0.288 | 0.96 | 0.556 | 1 | -0.323 | 0.925 | 22.3 | 6.9 | 7.2 | 63.7 |
| Thyasiridae | 0.125 | 0.749 | -0.264 | 0.953 | 0.225 | 0.91 | -0.339 | 0.948 | 14.4 | 9.9 | 27.3 | 48.4 |
| Lacernidae | 0.121 | 0.738 | 0.086 | 0.709 | -0.307 | 0.931 | -0.352 | 0.929 | 10.3 | 7.1 | 2.2 | 80.4 |
| Mopaliidae | 0.129 | 0.754 | 0.092 | 0.713 | 0.278 | 0.963 | -0.367 | 0.943 | 9.7 | 7.6 | 6.4 | 76.3 |
| Pleustidae | 0.015 | 0.531 | -0.091 | 0.718 | -0.232 | 0.864 | -0.39 | 0.953 | 11.1 | 10.7 | 14.2 | 64 |
| Oerstediidae | 0.176 | 0.807 | -0.167 | 0.847 | -0.195 | 0.815 | -0.418 | 0.955 | 12.3 | 10.8 | 8.2 | 68.7 |
| Entoprocta † | -0.347 | 0.953 | -0.103 | 0.741 | -0.015 | 0.533 | -0.43 | 0.974 | 9.8 | 8.4 | 13.1 | 68.7 |
| Cyamiidae | 0.108 | 0.706 | -0.117 | 0.763 | -0.213 | 0.842 | -0.437 | 0.967 | 12.9 | 13.8 | 15.8 | 57.4 |
| Spionidae | 0.099 | 0.693 | -0.22 | 0.911 | 0.279 | 0.958 | -0.466 | 0.977 | 7.8 | 8.4 | 5.5 | 78.3 |
| Mean | 0.088 | 0.754 | -0.076 | 0.738 | 0.154 | 0.846 | -0.013 | 0.81 | 13.494 | 5.535 | 10.931 | 70.043 |

**Supplementary Table 9:** MCMC convergence diagnostics for the three joint species distribution models in HMSC (six chains, 100,000 transient iterations, 4,000 samples per chain retained at a thinning of 100; 24,000 posterior samples in total). For each parameter group (Beta, fixed-effect coefficients; Omega, residual associations among taxa from the site random effect; Sigma, residual variance), 'ESS_min', 'ESS_median' and 'ESS_mean' summarise the effective sample size across parameters, and 'ESS_pct_gt200' gives the percentage of parameters with ESS > 200. 'PSRF_max' and 'PSRF_median' summarise the Gelman–Rubin potential scale reduction factor, and 'PSRF_pct_gt1.05' gives the percentage of parameters exceeding 1.05. Sigma is not estimated in the probit model, where the residual variance is fixed at 1.

| **Model** | Parameter | ESS_min | ESS_median | ESS_mean | ESS_pct_gt200 | PSRF_max | PSRF_median | PSRF_pct_gt1.05 |
|---|---|---|---|---|---|---|---|---|
| Subdivisions (CLR, normal) | Beta | 13977 | 23859 | 22782 | 100 | 1.001 | 1.000 | 0.000 |
| Subdivisions (CLR, normal) | Omega | 13279 | 22161 | 21067 | 100 | 1.007 | 1.000 | 0.000 |
| Subdivisions (CLR, normal) | Sigma | 17430 | 23937 | 23470 | 100 | 1.001 | 1.000 | 0.000 |
| Metazoan families (CLR, normal) | Beta | 8762 | 23931 | 23697 | 100 | 1.025 | 1.000 | 0.000 |
| Metazoan families (CLR, normal) | Omega | 4643 | 23364 | 22958 | 100 | 1.290 | 1.000 | 0.000 |
| Metazoan families (CLR, normal) | Sigma | 5513 | 24000 | 23508 | 100 | 1.001 | 1.000 | 0.000 |
| Metazoan families (presence–absence, probit) | Beta | 14296 | 23791 | 23204 | 100 | 1.001 | 1.000 | 0.000 |
| Metazoan families (presence–absence, probit) | Omega | 12132 | 23327 | 22532 | 100 | 1.065 | 1.000 | 0.000 |
| Metazoan families (presence–absence, probit) | Sigma | NA | NA | NA | NA | NA | NA | NA |
